# Integrative multi-omics landscape of non-structural protein 3 of severe acute respiratory syndrome coronaviruses

**DOI:** 10.1101/2021.07.19.452910

**Authors:** Ruona Shi, Zhenhuan Feng, Xiaofei Zhang

## Abstract

The coronavirus disease 2019 (COVID-19) caused by severe acute respiratory syndrome coronavirus 2 (SARS-CoV-2) infection is currently a global pandemic. Extensive investigations have been performed to study the clinical and cellular effects of SARS-CoV-2 infection. Mass spectrometry-based proteomics studies have revealed the cellular changes due to the infection and identified a plethora of interactors for all SARS-CoV-2 components, except for the longest non-structural protein 3 (NSP3). Here, we expressed the full-length NSP3 proteins of SARS-CoV and SARS-CoV-2 to investigate their unique and shared functions using multi-omics methods. We conducted interactome, phosphoproteome, ubiquitylome, transcriptome, and proteome analyses of NSP3-expressing cells. We found that NSP3 plays essential roles in cellular functions such as RNA metabolism and immune response such as NF-*κ*B signal transduction. Interestingly, we showed that SARS-CoV-2 NSP3 has both endoplasmic reticulum and mitochondrial localizations. In addition, SARS-CoV-2 NSP3 is more closely related to mitochondrial ribosomal proteins, whereas SARS-CoV NSP3 is related to the cytosolic ribosomal proteins. In summary, our multi-omics studies of NSP3 enhance our understanding of the functions of NSP3 and offer valuable insights for the development of anti-SARS strategies.

## Introduction

A few clusters of severe pneumonia from unknown sources were reported from Wuhan at the end of 2019 [1]. The causative factor of this pneumonia was soon isolated, sequenced, and designated as severe acute respiratory syndrome coronavirus 2 (SARS-CoV-2), and the disease was named coronavirus disease 2019 (COVID-19) [1–4]. Although approximately 80% of SARS-CoV-2-infected patients have mild to no symptoms, 15% of patients could develop pneumonia and dyspnea, whereas the remaining 5% of patients, especially those with chronic disease, including lung diseases or asthma, are at high risk of developing severe illness with high rates of mortality and morbidity [5, 6]. As SARS-CoV-2 is easily transmitted via respiratory droplets, COVID-19 has spread worldwide very rapidly and became a pandemic by March 2020 [7]. As of 19th, July 2021, more than 191 million infections and 4 million deaths were reported according to the Center for Systems Science and Engineering at the Johns Hopkins University (https://coronavirus.jhu.edu/). Encouragingly, the research society has made tremendous progress in understanding of the molecular mechanisms behind the viral infection, as well as in the development of antiviral compounds and vaccines to treat COVID-19 [8, 9].

SARS-CoV-2 is a beta coronavirus and possesses a ∼30 kb positive, single-strand genome. Phylogenetic sequence analysis revealed that the SARS-CoV-2 is closely related to two human transmissible beta coronaviruses, SARS-CoV (80% gene sequence similarity) and MERS-CoV (50% gene sequence similarity) [6, 10]. Bioinformatic predictions suggest that SARS-CoV-2 encodes four structural proteins, nine accessory proteins, and two long polypeptides (pp1a and pp1ab) [4, 11]. These two long polypeptides are translated from *ORF1a* and *ORF1ab*. *ORF1ab* is produced via a −1 ribosomal frameshift from the stop codon of *ORF1a*, and these two polypeptides together produce a total of 15 non-structural proteins (NSPs) via proteolysis. The processing of pp1a and pp1ab requires the protease activity of NSP3 (papain-like protease) and NSP5 (chymotrypsin-like protease, 3CLPro) [11].

Previous studies on SARS-CoV revealed that the NSPs are the primary constituents for the assembly of the replication and transcription complex (RTC), where viral genome RNA synthesis occurs and dsRNA is abundantly expressed [12]. Co-expression of NSP3, NSP4, and NSP6 induce the formation of double-membrane vesicles (DMVs), which are continuous with the endoplasmic reticulum (ER) membrane. These DMVs have similar morphology to the organelle-like structures induced by SARS-CoV infection [12, 13]. These multiple proteins and membrane containing organelle-like RTC facilitates the synthesis of positive-strand RNA viruses where no interference from the host cell innate immune system occurs [14]. The NSP3 of SARS-CoV is the largest and most essential component of the RTC complex. It has 1922 amino acids and multiple functional domains with ssRNA binding, deMARylation, G-quadruplex binding, and cysteine protease activity [15, 16]. Therefore, it is not surprising that NSP3 plays pivotal roles in the viral life cycle, and the inhibition of NSP3 PLpro activity prevents SARS-CoV virus replication [17]. NSP3 of SARS-CoV-2 is a 1945 amino acids polypeptide that has conserved functional domains with an overall 86% protein sequence similarity to that of SARS-CoV. Recent studies have demonstrated that the deMARylation, deubiquitination and deISGylation functions of the Macro and PLpro domains of SARS-CoV-2 [16, 18–20]. More importantly, two studies showed that PLpro inhibition results in excellent anti-SARS-CoV-2 activity [18, 19]. However, despite years of extensive research into the functions of NSP3 from SARS-CoV and recently into SARS-CoV-2, the functions of full-length NSP3 are still only partially understood. We only found two reports that successfully expressed full-length NSP3, one for SARS-CoV, and one for SARS-CoV-2 [12, 19]. In addition, three very recent interaction studies were performed for SARS-CoV-2 NSP3, with two studies using deletion mutants, whereas the third one failed to detect the expression of NSP3 [21–23].

In this study, we successfully cloned and expressed NSP3 protein of SARS-CoV and SARS-CoV-2. Using these plasmids, we present the transcriptomic and proteomic (interactome, proteome, phosphoproteome and ubiquitylome) landscapes of NSP3. These parallel studies were used to investigate the similarities and differences between NSP3 proteins. Specifically, we investigated how NSP3 protein interacts with host cells. Furthermore, we assessed the signaling pathways that are regulated by the NSP3 protein. Lastly, we also aimed to offer a list of compounds that could potentially be repurposed for further investigations of COVID-19 treatment.

## RESULTS

### Expression of NSP3 proteins

Current studies into the functions of NSP3 proteins (Figure 1A) of both SARS-CoVs (hereafter referred to as CoV1-NSP3 and CoV2-NSP3) are mainly limited to their truncated deletions. We successfully cloned the full-length, wild-type (WT), catalytically inactive (CS), or dGG-mutated (last two glycine residues deleted, resistant to PLpro cleavage), NSP3 proteins of both SARS-CoVs (without codon optimization) into a lentivirus vector, as illustrated in Figure S1A. We expressed all six plasmids in HEK293T cells and detected their expression using EGFP and CoV1-NSP3 antibodies. As shown in Figure S1B, the expression of CS- and dGG-mutated NSP3 proteins was detected using an EGFP antibody (∼250 kDa), whereas no obvious expression of WT-NSP3 was detected. This is due to the self-cleavage activity of PLpro toward the Leu-Lys-Gly-Gly of its C-terminal, as dGG-NSP3 is expressed well [24]. We verified this hypothesis by treating cells with the PLpro inhibitor GRL0617 [17, 18]. As shown in Figure 1B, EGFP antibody could detect the WT-NSP3 proteins in GRL0617-treated cells. This observation also explains why a previous study failed to detect the expression of NSP3 using a C-terminal tag [22].

**Figure 1.**
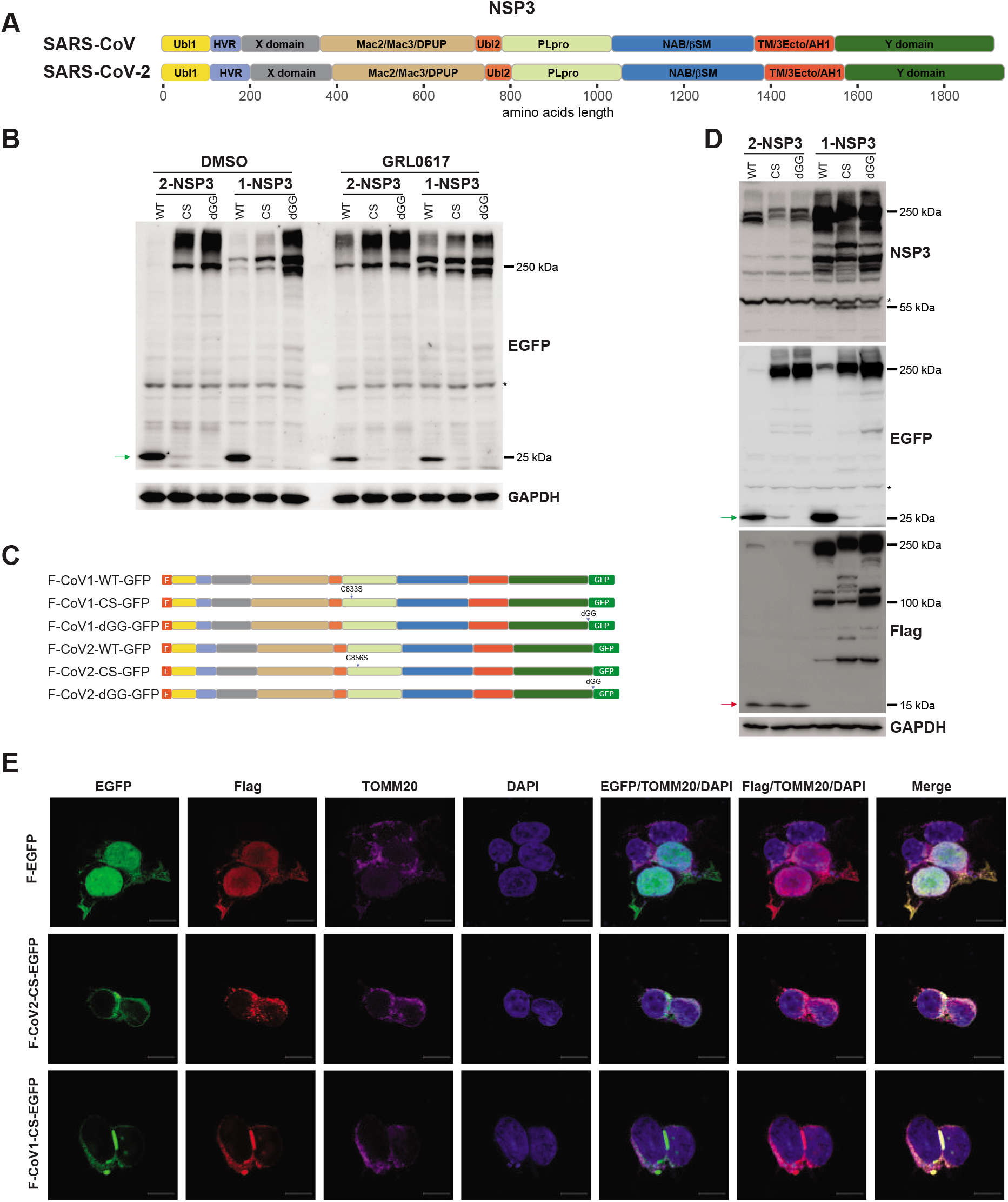
Expression and subcellular localization of SARS-CoVs NSP3 proteins. **A.** Schematic illustration of SARS-CoVs NSP3 proteins. Domain illustration of NSP3 proteins of SARS-CoV and SARS-CoV-2, respectively [15, 16]. Ubl1 is ubiquitin like domain 1, HVR is hyper variable region, X/Mac1 is macrodomain I, Mac2/Mac3/DPUP are macrodomain II, III and domain preceding Ubl-2 and PLpro, Ubl2 is ubiquitin like domain 2, PLpro is Papain-like protease 2, NAB/βSM is nucleic acid-binding and beta-coronavirus-specific marker domain, TM1 is transmembrane region 1, 3Ecto is 3 ectodomain, TM2 is transmembrane region 2, AH1 is amphipathic helix1, Y domain. **B.** GRL0617 inhibits the release of EGFP from NSP3-EGFP protein. Cells transfected with indicated plasmids (Figure S1A) were treated with DMSO or GRL0617 for 24 h. Cells were lysed and EGFP antibody was used for immunoblotting. Green arrow indicates released EGFP protein, and asterisk indicates non-specific signal. GAPDH was used to serve as a loading control. **C.** Schematic illustration of plasmids used in Figure 1D. **D.** CoV2-NSP3 is processed at the N-terminus. Cells transfected with indicated plasmids (Figure 1C) were analyzed as in Figure 1B. Green arrow indicates the released EGFP and red arrow around 15 kDa indicates processed N-terminus of CoV2-NSP3. Asterisks indicate non-specific signals. **E.** N-terminus of CoV2-NSP3 prefers to co-localize with TOMM20. HEK293T cells transfected with Flag-CoVs-NSP3-EGFP plasmids were fixed 48 hr. post-transfection. Flag and TOMM20 antibodies were used to visualize the co-localization.

Next, we performed immunoblotting using the NSP3 antibody which is specific for CoV1-NSP3. The specificity of this CoV1-NSP3 antibody to CoV2-NSP3 was verified by immunoblotting and immunostaining (Figure S1B and S1C). Interestingly, we noticed that CoV2-NSP3 had a lower molecular weight than CoV1-NSP3, with both EGFP and NSP3 antibodies (Figure 1B and Figure S1B). This implies that either the amino acid composition of NSP3 protein affects its molecular weight or that CoV2-NSP3 undergoes extra processing at its N-terminus. We therefore subcloned NSP3 into a vector with N-terminal Flag and C-terminal EGFP tags (Figure 1C). Interestingly, we observed a very specific processed band of approximately 15 kDa for all three CoV2-NSP3 proteins (Figure 1D, Flag antibody). In addition, the Flag antibody detected most of CoV1-NSP3 at the correct position, thought multiple smaller signals were also observed, suggesting the existence of extra processing (Figure 1D).

### Localization of NSP3 proteins

We sought to determine the cellular localization of the NSP3 protein by immunostaining. As shown in Figure S1C, EGFP and NSP3 signals were well-localized, especially in CS- and dGG-NSP3-expressing cells. Interestingly, cells expressing CS-CoV1-NSP3 showed very large vesicle-like structures, which were not observed with WT and dGG-mutated proteins. Next, we stained NSP3-expressing cells with ER, autophagosome, endosome, and mitochondrial makers, as all four organelles are potential origins of DMVs [25–28]. Both NSP3 proteins showed no obvious co-localization with the endosome marker RAB7 or autophagosome marker LC3 (data not shown). As shown in Figure S2A, we found that CS-CoV1-NSP3 co-localized with the ER marker calnexin (CANX), particularly in the large vesicle-like structures, but not with TOMM20 (Figure S2B). In contrast, the CoV2-NSP3 showed partial co-localization with both TOMM20 and CANX (Figure S2A and S2B). We next investigated whether the extra N-terminal processed band might somehow affect the localization of CoV2-NSP3. As shown in Figure S2C, NSP3 antibody showed fewer co-localization signals with Flag-CoV2-NSP3, compared to the co-localization signals of NSP3 and EGFP antibodies. In addition, the Flag antibody also detected vesicle-like structures, though smaller in size than those in CoV1-NSP3-expressing cells (Figure S2C). These vesicle-like structures showed strong co-localization with TOMM20, but not with CANX (Figure 1E and S2D). Interestingly, the EGFP signal of CoV2-NSP3 still co-localized with CANX (Figure S2D). These observations indicated that there are two cellular localizations of CoV2-NSP3, influenced by the extra processing at its N-terminus. In contrast, Flag staining of CoV1-NSP3 showed the same localization pattern with CANX as with EGFP (Figure S2D). Finally, extracting data from interactome studies verified the localization preference of different NSP3 proteins (Figure S2E).

### The N-terminus determines the localization of CoV2-NSP3

To further investigate the cellular localization of NSP3 proteins, we exchanged their Ubl1 and HVR domains reciprocally (Figure 2A). We also exchanged the PLpro domain as the CS-CoV1-NSP3 mutant shows large vesicle-like structures. As shown in Figure 2B, when the Ubl1 and HVR domains of CoV2-NSP3 were changed to those of CoV1-NSP3, the N-terminal extra band was disappeared. In contrast, a 15 kDa band was observed when CoV1-NSP3 was equipped with the Ubl1 and HVR domains of CoV2-NSP3. In addition, we showed that the PLpro domain has no obvious influence on processing of the CoV2-NSP3 protein (Figure 2B).

**Figure 2.**
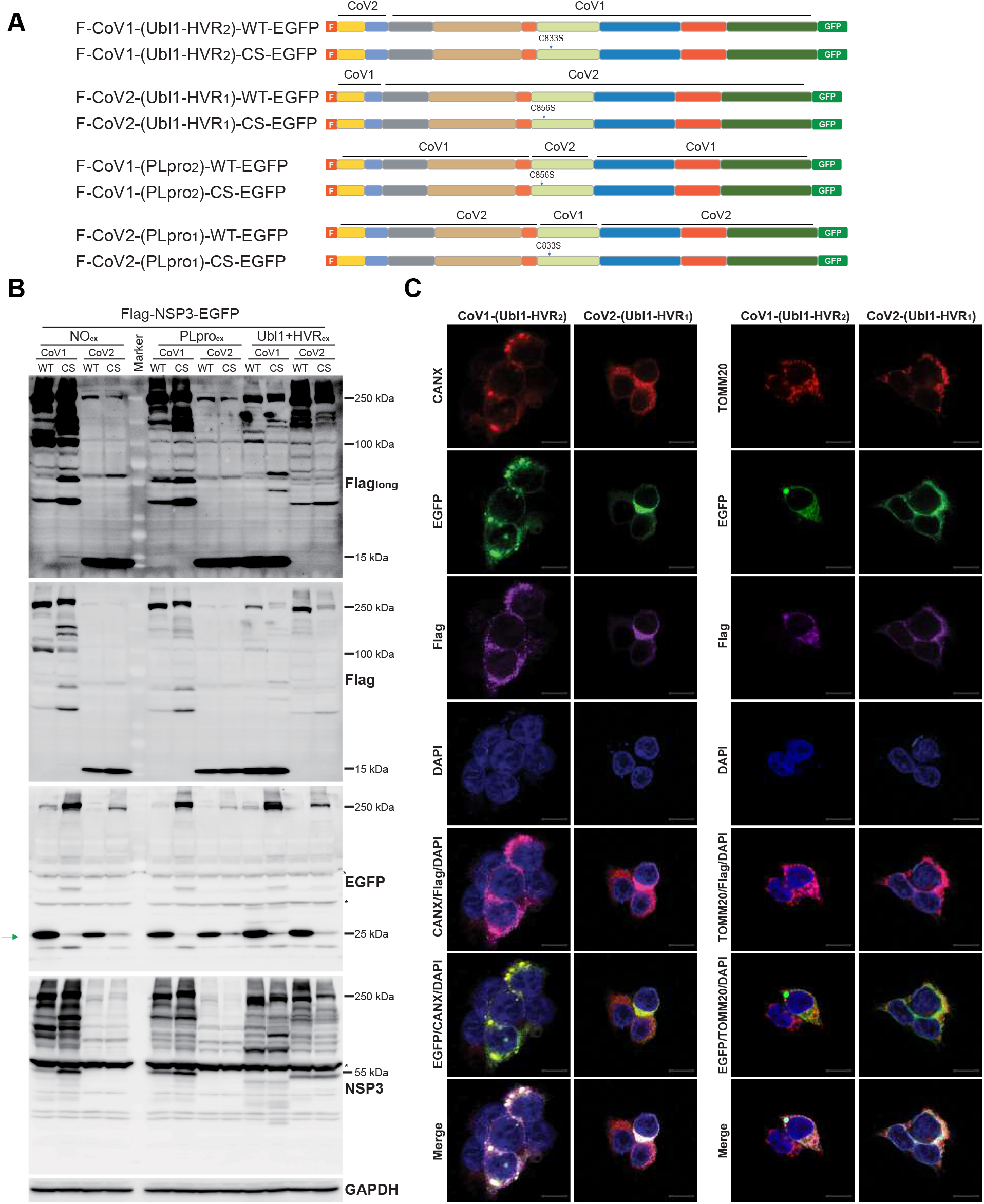
N-terminus determines the extra processing and mitochondria localization of NSP3. **A.** Schematic illustration of chimeric plasmids used in Figure 2. **B.** The processing site of CoV2-NSP3 is localized at its N-terminus. HEK293T cells transfected with indicated plasmids were analyzed as in Figure 1B. Green arrow indicates release EGFP. PLpro_ex_ indicates the exchange of PLpro domain, whereas Ubl1+HVR_ex_ indicates the exchange of Ubl1 and HVR domains. Asterisks indicate non-specific signals. **C.** The N-terminus of CoV2-NSP3 re-locates CoV1-NSP3 to mitochondria. HEK293T cells transfected with indicated plasmids were analyzed as in Figure 1E.

Next, we used immunostaining to study the localization of these fused proteins (Figure 2C). As expected, CoV1-NSP3, detected with the Flag antibody, co-localized with TOMM20 when its N-terminus was replaced to that of CoV2-NSP3. In contrast, CoV2-NSP3 fused with the N-terminus of CoV1-NSP3 was predominantly co-localized with CANX. However, the EGFP signals of both fused proteins was still co-localized with CANX. Therefore, we concluded that the C-terminus of both NSP3 proteins co-localizes with ER, whereas the N-terminus of CoV2-NSP3 prefers to bind to mitochondria. Furthermore, we showed that the exchange of the PLpro domain has no obvious influence on NSP3 protein localization (Figure S3A).

### Interactome analysis of NSP3

To study how NSP3 protein contributes to cellular function regulation during infection, we conducted a multi-omics study (interactome, phosphoproteome, ubiquitylome, transcriptome) on NSP3-expressing cells (Figure S4A). We performed all experiments in biological triplicates, and we showed that the distribution of total intensity was consistent within every experimental condition (Figure S4B). We first analyzed the interactome of NSP3 using Flag-immunoprecipitation coupled with mass spectrometry identification. As shown in Figure S5A, principal component analysis (PCA) of the top 1000 variable proteins showed a strong correlation within the triplicate experiments for both NSP3 proteins. In addition, the interactome of the WT and CS mutants of NSP3 showed a good correlation, implying that the catalytic activity has minor effects on the NSP3 interactome. In total, 221 and 226 of significant interactors were enriched for CoV1-NSP3 and CoV2-NSP3, respectively, using thresholds based on the fold-change (> 4) and an adjusted *P* < 0.001. Next, we compared these two NSP3 interactomes side by side to study their shared and specific interactions (Figure 3A). We noticed that RCHY1, a known interactor of the macrodomain and PLpro of NSP3, was enriched with both NSP3 proteins [29]. However, the overall similarities between the two NSP3 interactomes was low (∼15%, Figure 3A and Figure S5B). Gene ontology (GO) and Reactome [30] analyses of significant interactors highlighted the roles of both NSP3 proteins in RNA splicing, NF-*κ*B signaling, and the regulation of translation (Figure 3B, Figure S5C, S5D). The most striking difference between the two interactomes was that CoV2-NSP3 was linked to mitochondrial translation, whereas CoV1-NSP3 was associated with cytosolic translation. We verified a few specific and shared interactions using immunoprecipitation followed by immunoblotting (Figure 3C). For example, TAB2 and TAB3 showed similar binding ability to both NSP3 proteins, whereas cytosolic ribosomal proteins and mitochondria ribosomal proteins preferred to bind to CoV1-NSP3 and CoV2-NSP3, respectively. To avoid missing of transient protein-protein interactions and to identify a core protein-protein interaction complex, we repeated our interactome study using a crosslinking method (Figure S6A-S6D) called rapid immunoprecipitation mass spectrometry of endogenous proteins [31]. Approximately 30% and 55% of high confident interactors (Figure S6B) from crosslinking experiments were found in non-crosslinking experiments for CoV2-NSP3 and CoV1-NSP3, respectively. Strikingly, most of the interactors of CoV2-NSP3 were mitochondrial proteins including mitochondrial ribosomal protein large subunits (MRPLs), mitochondrial ribosomal protein small subunits (MRPSs), and coenzyme Q (COQs) (Figure S6C). In addition, biological process (BP) analysis showed that proteins that related to the mitochondrial gene expression, quinone biosynthetic process, and cellular respiration were significantly enriched with CoV2-NSP3 (Figure S6D), further validating the intimate interaction between CoV2-NSP3 and mitochondria.

**Figure 3.**
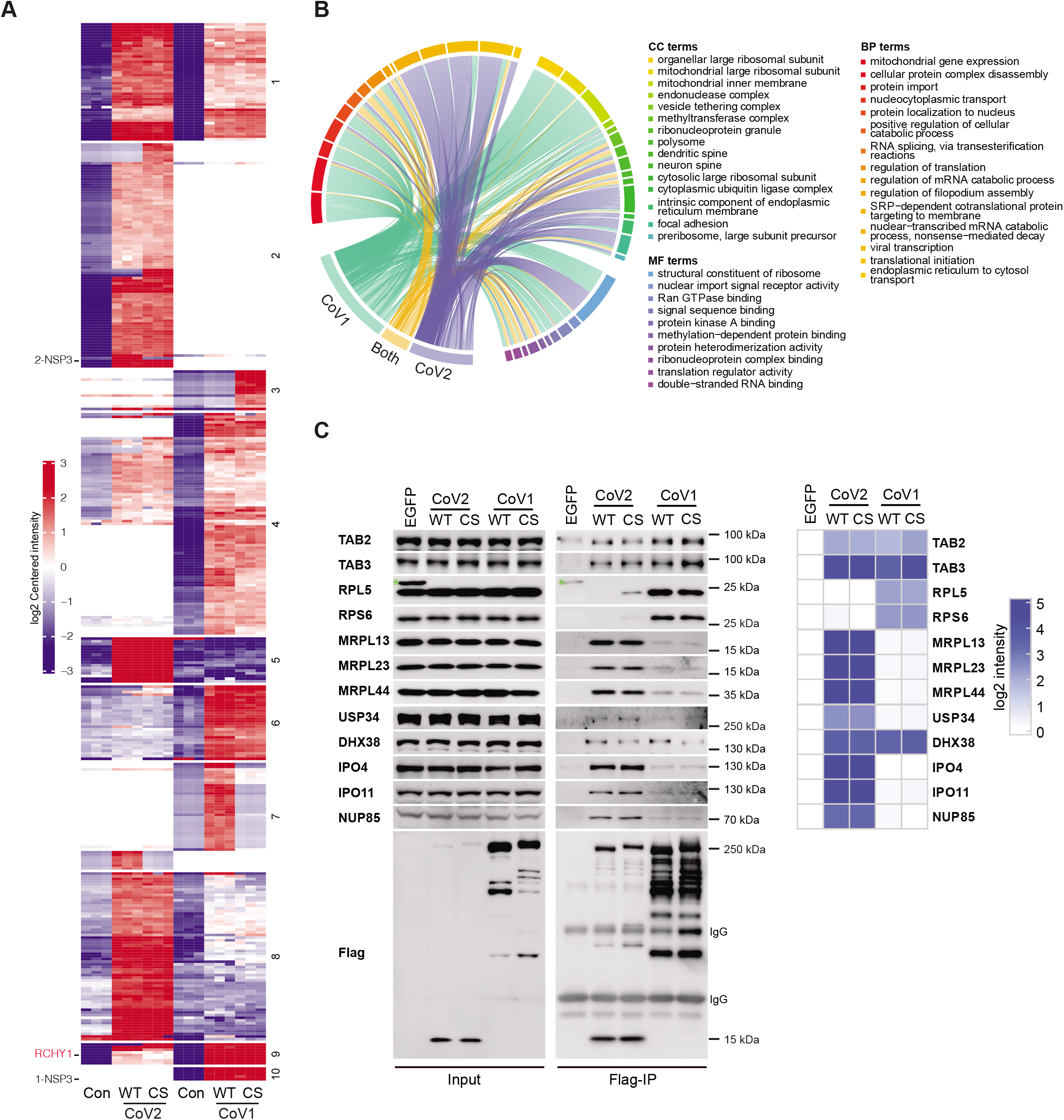
Interactome analysis of NSP3 proteins. **A.** Clustering of significant interactors of NSP3 proteins. Significant interactors of NSP3 were analyzed to show the shared and unique interactions. Data analysis as described in material and methods. Red indicates enrichment, while blue indicates lack of enrichment. Interactors that do not present in either NSP3 interactome are left empty. RCHY1 and NSP3 proteins are indicated in left. **B.** Representative enriched GO of identified interactors. GO analysis was performed as described in material and method. Circos diagram was used to show the representative enriched GO terms. BP represents biological process; CC represents cellular component and MF represents molecular function. **C.** Validation of interactome study. HEK293T cells transfected with Flag tagged NSP3 plasmids were harvest for Flag immunoprecipitation. Indicated antibodies were used to detect the interactions. LFQ intensity of each interactor was log2 transformed and then substrate its intensity identified in EGFP condition. Black asterisk indicates non-specific signal, whereas green asterisk indicates leaked EGFP signal.

Because the extra processing of CoV2-NSP3 at its N-terminus, it was therefore necessary to study the interactome with a tag localized to the C-terminus of the NSP3 protein (Figure S7A-S7D). As shown in Figure S7B, RCHY1 was again enriched with both NSP3 proteins again. The overall similarities between the two EGFP interactomes was a slightly higher than those of Flag-IP (27% vs. 15%, Figure S5B and Figure S7C). Consistent with immunostaining results, interactors of both NSP3 proteins were related to the ER (Figure S7D). In addition, shared interactors were enriched in IL-1 signaling. We next compared our result with a recent published NSP3 interactome using deletions [23]. As shown in Figure S7E, approximately 20% of our interactors were also found in NSP3 deletions study.

### Viral protein-host protein interaction network

To study how the NSP3 protein is connected to other SARS-CoV-2 proteins and host proteins, we mined our data with reported SARS-CoV-2 interactors [29, 32]. Here, we combined all of our identified NSP3 interactors with reported interactors for all other SARS-CoV-2 proteins. As shown in Figure 4, NSP3 was linked to multiple viral proteins via its interacting partners, suggesting that they work coordinately to regulate host cell functions. We found that NSP3, ORF3, and NSP6 shared multiple ER-localized interactors, which is consistent with the observation of ER-localization of both NSP3 proteins. Furthermore, both NSP3 proteins were linked to NSP6 via ATP metabolism-related proteins. Interestingly, only CoV2-NSP3 was associated with NSP8 and M proteins via mitochondrial-related proteins. In addition, ORF7A exclusively interacted with CoV2-NSP3, but not CoV1-NSP3. These results indicated that NSP3 coordinates with other viral proteins to regulate cellular functions.

**Figure 4.**
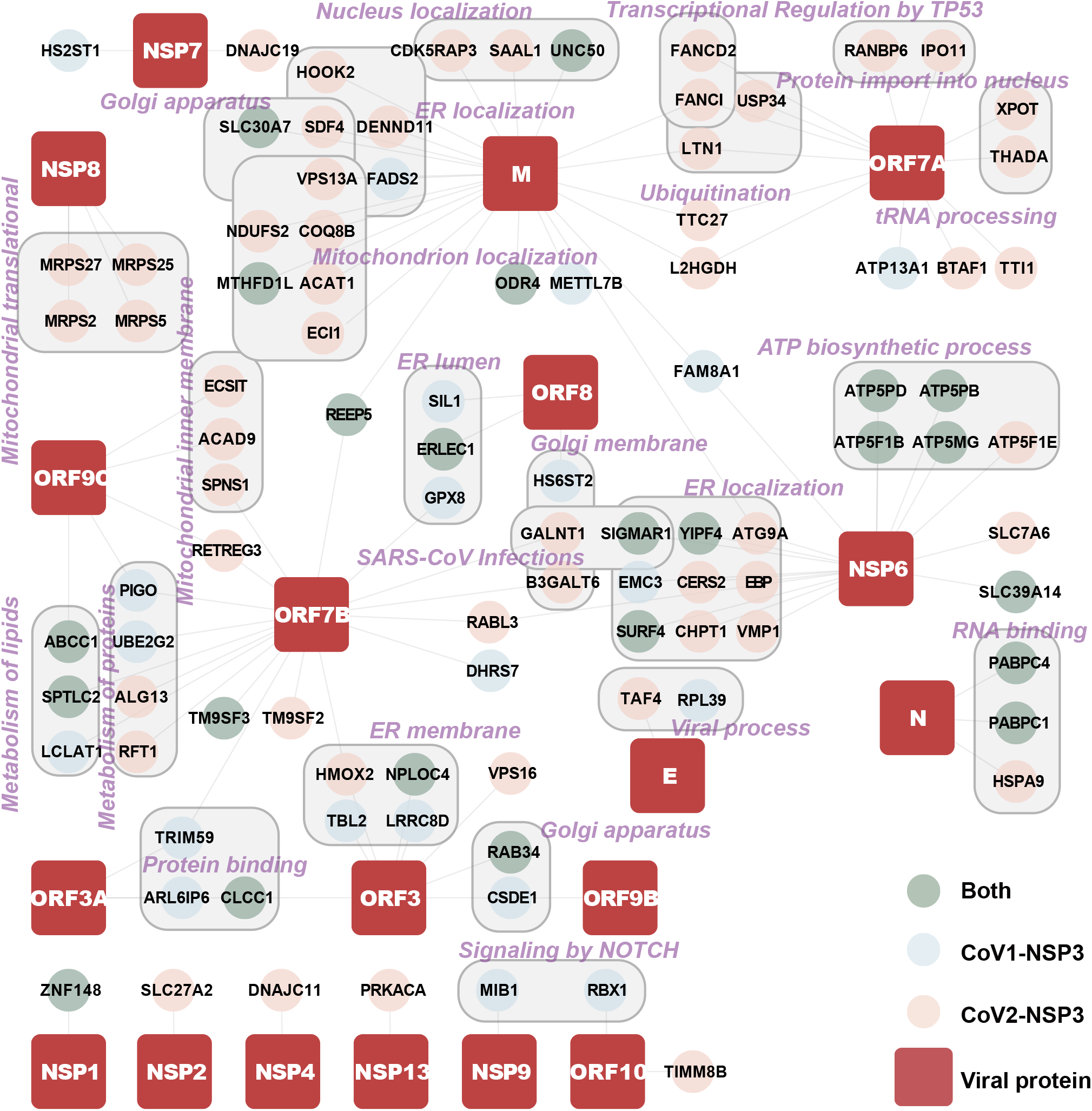
Viral protein-host protein interaction network. **A.** Interaction connections of NSP3 with other SARS-CoV-2 proteins. The interactors of NSP3 proteins were combined with reported SARS-CoV-2 interactors to create viral protein-host protein interaction network. Interactors specific for CoV1-NSP3 and CoV2-NSP3 are displayed as different colors. Cytoscape was used to create the connection map of NSP3, interactors and SARS-CoV-2 proteins as described in material and methods.

Taken together, our interactome results indicated the following: 1) regulation of RNA metabolism could be the main function of NSP3; 2) both NSP3 proteins are associated with protein translation but employ different translation machineries; 3) The N-terminus of CoV2-NSP3 determines the interaction with mitochondria; and 4) NSP3 functions together with other SARS-CoV-2 proteins to control host cell functions.

### Phosphoproteome analysis of NSP3-expressing cells

We next explored the effects of NSP3 on the phosphoproteome of host cells using a highly sensitive phosphoproteome preparation and identification workflow (EasyPhos) [33]. Different enriched phosphorylated peptides (DEPPs) were identified if they appeared at least three times in one condition based on an ANOVA test, with thresholds of a Benjamini-Hochberg adjusted to *P* < 0.01, and |L2FC| > 2. For all analyses, we chose 0.7 as the minimum cutoff for the localization probability. In total, 659 DEPPs were identified for both NSP3-expressing cell lines. Moreover, some of the DEPPs showed specificity in either NSP3-expressing cell line (Figure 5A). We validated the serine and threonine phosphorylation of UBE2O and TNIP1 using immunoprecipitation followed by immunoblotting (Figure 5B). BP and Reactome analyses of NSP3 shared DEPPs (Figure 5A and Figure 5C, cluster 5 and 6) showed that terms related RNA metabolism, the regulation of viral transcription, and potential therapeutics for SARS were enriched with both NSP3 proteins. Interestingly, similar BP and Reactome terms were found for DEPPs in cluster 2 and cluster 4, although they were specific DEPPs for CoV2-NSP2 and CoV1-NSP3, respectively. To dissect the contributions of NSP3 to the host cell phosphoproteome during viral infection, we compared our results with recently reported SARS-CoV-2 phosphoproteome datasets [34, 35]. In total, 90 DEPPs in our dataset were matched to previous reports, and a large proportion of them showed similar regulation trends (Figure S8B). For example, DEPPs with similar regulatory trends were found for SRRM2, UBE2O, SRSF3 and SIPA1L1, emphasizing the contributions of NSP3 to these phosphorylation events during viral infection.

**Figure 5.**
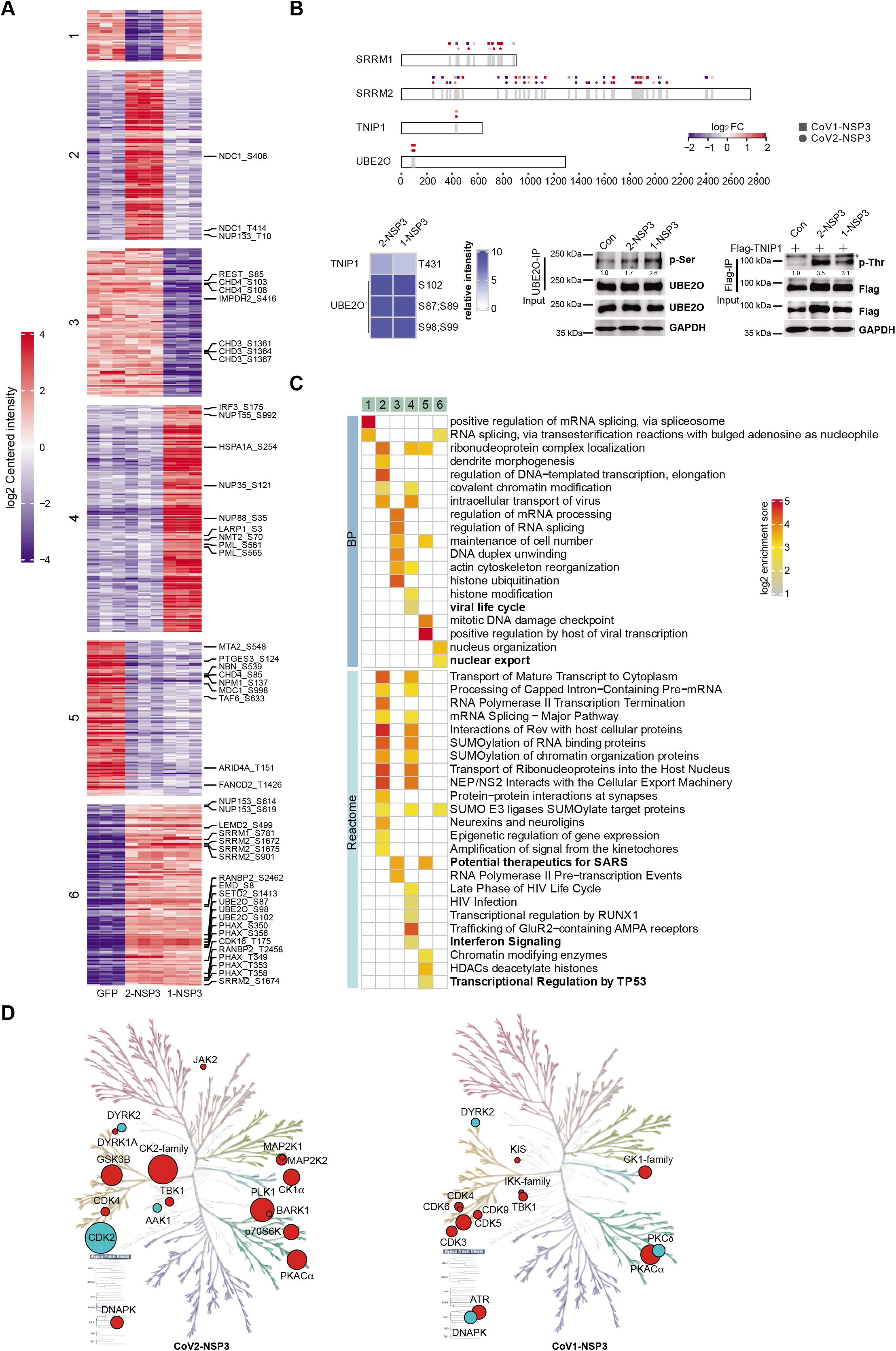
Phosphoproteome analysis of NSP3-expressing cells. **A.** Clustering of DEPPs in NSP3-expressing cells. DEPPs show significantly differences in CoV1-NSP3- and CoV2-NSP3-expressing cells compared to control cells were clustered together. Analysis and colors as in Figure 3A. Representative DEPPs with highlighted GO in Figure 5C were shown on the right. **B.** Validation of phosphoproteome studies. Top panel: Schematic representation of phosphorylated sites of indicated proteins. Square indicates DEPP has significant change in CoV1-NSP3-expressing cells, whereas circle indicates DEPP has significant change in CoV2-NSP3-expressing cells. Analysis and colors as in Figure 3A; Bottom panels: HEK293T cells transfected with indicated plasmids were subjected to immunoprecipitation (Flag-resin or UBE2O antibody with protein A/G), following immunoblotting with phosphorylated serine, threonine antibodies. ImageJ was used to quantify the relative expression of each protein to its matching loading control. **C.** Representative enriched biological process (BP) and Reactome. Label on top of the heatmap corresponds to the cluster in Figure 5A. Bold terms containing DEPPs are highlighted in the right of Figure 5A. **D.** Kinmaps show altered kinases whose activity mapped to DEPPs in NSP3-expressing cells. Significantly different enriched DEPPs in CoV1-NSP3- and CoV2-NSP3-expressing cells were matched to kinases. Kinome profiling was performed using KinMap interface. Red indicates upregulation, whereas blue indicates downregulation. Size of the circles reflects the number of DEPPs.

We then mapped our DEPPs to kinases that regulate these phosphorylation sites and visualized the data using KinMap (Figure 5D) [36]. We found that the altered activity of CDK4, CK1*α*, TBK1, and DYRK2 was conserved in both NSP3-expressing cell lines. However, the activities of multiple CDKs were upregulated in CoV1-NSP3-expressing cells, whereas only CDK4 activity was upregulated in CoV2-NSP3-expressing cells. The decrease in CDK2 activity in CoV2-NSP3-expressing cells is consistent with the results obtained from virus-infected cells [35]. As CDKs play an important role in cell cycle progression, we showed that the expression of both NSP3 proteins caused a prolonged G2 phase compared to that of control cells (Figure S8C). We also mapped inhibitors to kinases for which the activity was significantly changed in NSP3-expressing cells (Figure S8D and S8E). We retrieved 12 kinases and 18 kinase inhibitors from our phosphoproteome data of CoV2-NSP3-expressing cells (Figure S4D). Further, one of the inhibitors, fostamatinib, a treatment for chronic immune thrombocytopenia targeting multiple mapped kinase pathways [37], has been investigated as a potential treatment for acute lung injury for COVID-19 patients [38]. Interestingly, we noticed that the interferon activators, IKK-family and TBK1 kinases were significantly present in CoV1-NSP3-expressing cells (Figure S8E and S8F), indicating the different effects of NSP3 proteins.

### Ubiquitylome analysis of NSP3-expressing cells

Ubiquitination is used by both viruses and host cells to combat each other [39]. For SARS-CoVs, they both have a PLpro protease domain within the NSP3 that is involved in deubiquitination and deISGylation activities [18]. We used the antibody-based K-*ε*-GG peptide enrichment method to study the effects of WT and CS-mutated NSP3 on the ubiquitylome of the host cells. Because the levels of ISGylation and NEDDylation are relatively low in cells, we therefore refer to the enriched peptides as ubiquitinated peptides (ubiquitylome) hereafter.

In total, 449 differentially enriched ubiquitinated peptides (DEUPs) showed significant differences with thresholds of localization probability > 0.7, |L2FC| > 2, and adjusted *P* < 0.001 (Figure 6A, Figure S9A). We noticed that most of the ubiquitination events, except some in cluster 1, 4, and 8, were not consistent in NSP3-expressing cells. However, GO analysis showed similar enriched terms, such as positive regulation of viral life cycle, mRNA splicing, cell cycle regulation and translation initiation, for both NSP3 proteins (Figure 6B). These results indicated that both NSP3 proteins have similar cellular functions, although they might function through different substrates. In addition, only 22 DEUPs and 25 DEUPs showed catalytic activity-dependent regulation in CoV2-NSP3 and CoV1-NSP3-expressing cells, respectively (Figure S9B). This observation suggested that the marked effects of NSP3 on the cellular ubiquitylome is catalytic activity-independent, which was also observed for other deubiquitinases [40].

**Figure 6.**
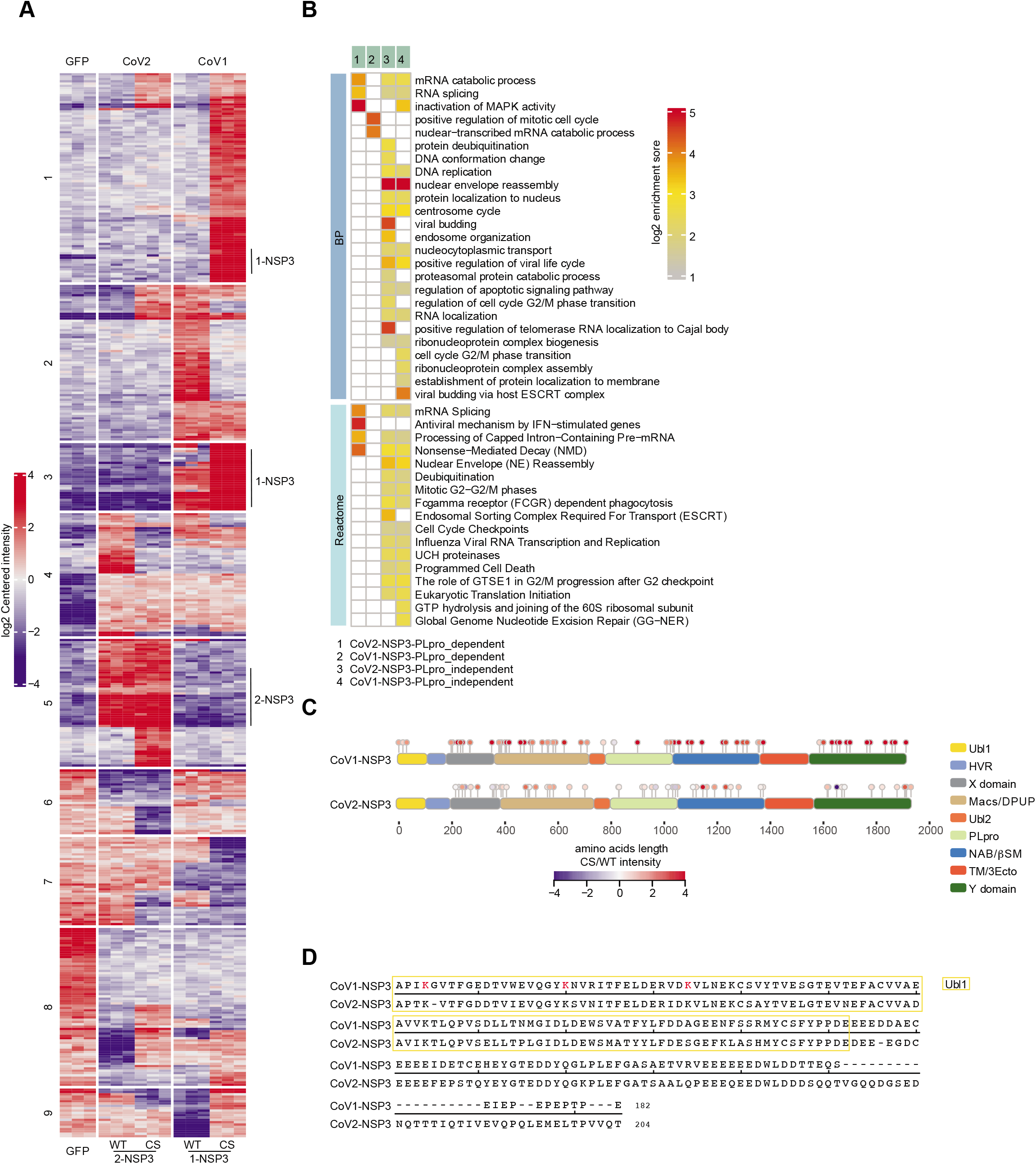
Ubiquitylome analysis of NSP3-expressing cells. **A.** Clustering of DEUPs in NSP3-expressing cells. DEUPs show significantly differences in NSP3-expressing cells compared to control cells were clustered together. Analysis and colors as in Figure 3A. **B.** BP and Reactome analyses of catalytic activity dependent- and independent-DEUPs. **C.** Schematic representation of identified ubiquitination sites of NSP3 proteins. The intensity of each ubiquitinated site was log2 transformed and compared between CS- and WT-mutated NSP3-expressing cells. Colors as in Figure 3A. **D.** Alignment of the Ubl1 and HVR domains of NSP3 proteins. Red-labelled K indicates identified ubiquitinated lysine residues of CoV1-NSP3.

We next mapped all ubiquitinate sites of NSP3 proteins to their functional domains and found that the overall ubiquitination sites were conserved in both NSP3 proteins. The three domains, HVR, Ubl2, and TM/3Ecto, had almost no ubiquitinated sites, explained by the fact that these domains only have a few lysine residues. Interestingly, we noticed that the N-terminus of CoV1-NSP3 had three ubiquitinated sites, whereas no such ubiquitination was observed in CoV2-NSP3 (Figure 6C). However, these three lysine amino acids were also conserved in CoV2-NSP3 (Figure 6D), raising questions about why the N-terminus of NSP3 proteins is differently ubiquitinated, and as well as the mechanisms underlying and consequences of these differences.

### Transcriptome and proteome changes in NSP3-expressing cells

We also measured the transcriptome and proteome of control and NSP3-expressing cells using deep-sequencing and filter-aided sample preparation (FASP)-based proteomics analyses. As shown in Figure 7A, differentially expressed genes (DEGs) were calculated using thresholds of an adjusted FDR < 0.05 and |L2FC| > 1 compared with the expression in control cells. Using motif enrichment analysis, we found that RELB-, NF-*κ*B1-, and NF-*κ*B2-binding motifs were highly represented in the upregulated DEGs, whereas ZBTB7A-, ZBTB7B- and AFF4-binding motifs were enriched in the downregulated DEGs (Figure 7A). We then matched the top 1000 variable genes of our gene sets with C2 (chemical and genetic perturbations sub-collection) and C5 (GO sub-collection) collections from the Molecular Signature Database (MsigDB) [41, 42]. As shown in Figure 7B, our upregulated genes were correlated with multiple reported gene sets, and a large portion of them was related to perturbations in NF-*κ*B signaling. More importantly, our gene sets had a significant correlation with that of SARS-CoV-2 infection of Calu-3 and A549 cells and respiratory syncytial virus infection of A549 cells.

**Figure 7.**
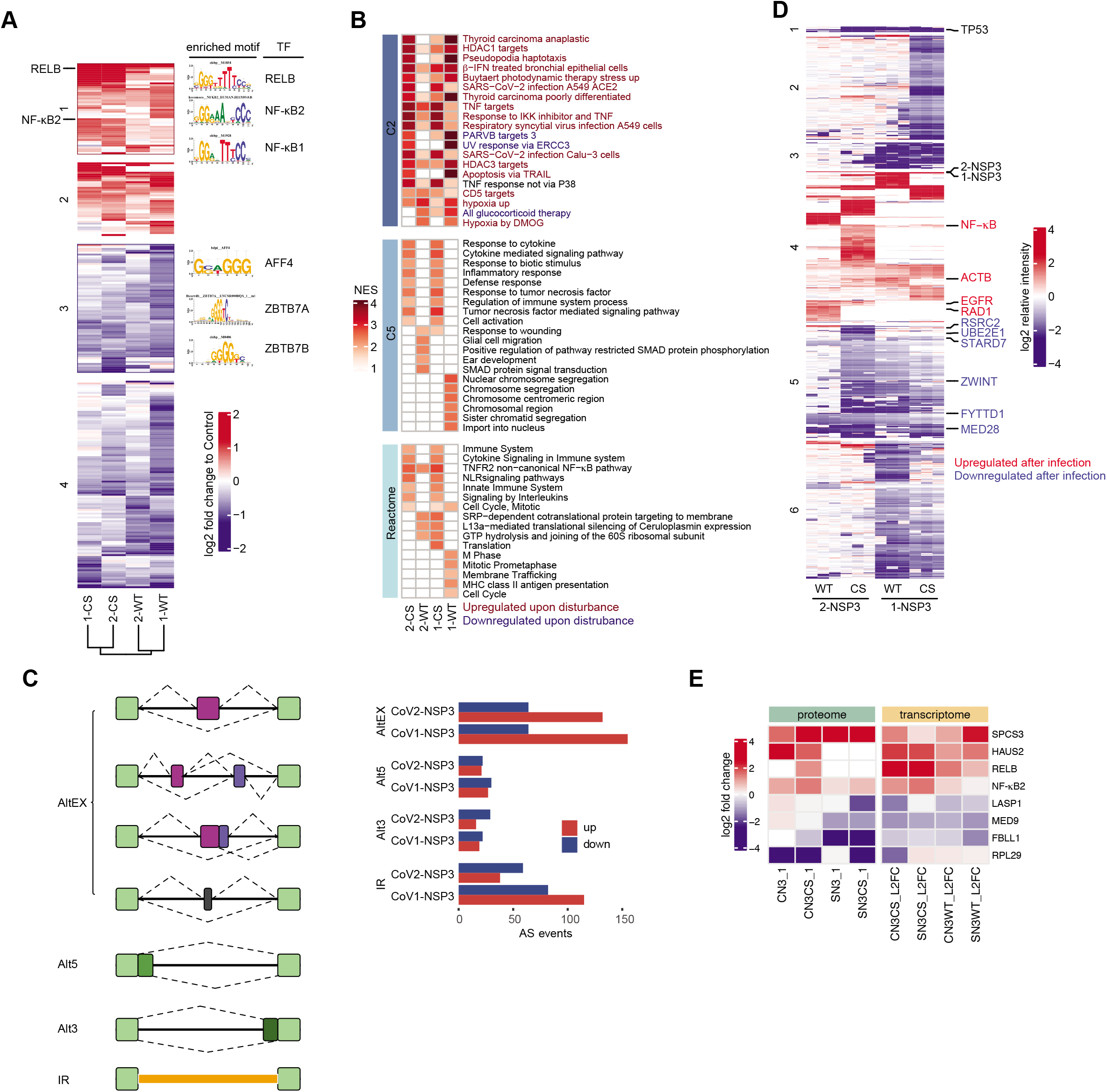
Transcriptome and proteome analyses of NSP3-expressing cells. **A.** Motif enrichment analysis of DEGs in NSP3-expressing cells. DEGs show significantly differences in NSP3-expressing cells were clustered together. Analysis and colors as in Figure 3A. Motif enrichment were performed as described in material and methods. RELB and NF-*κ*B were upregulated in NSP3-expressing cells. Enriched motifs are shown in the right of the heatmap. **B.** Gene set enrichment analyses of DEGs. C2 (curated gene sets) and C5 (GO gene sets) from MSigDbv7.2 and pathway gene sets from Reactome pathway database were used for GSEA. NES represents negative normalized enrichment score. **C.** Alternative splicing events in NSP3-expressing cells. Schematic illustration of alternative splicing events identified in NSP3 expressing cells (left panel). AS, alternative splicing events identified from the RNA-seq data of NSP3-expressing cells compared to control cells. AltEX represents alternative exon, Alt5 and Alt3 represents alternative 5’ and 3’ UTR, IR represents intron retention. **D.** Hierarchical clustering of DEPs in NSP3-expressing cells. Analysis and colors as in Figure 3A. Proteins showed consistent regulation in viral infection are labelled in the right of the heatmap [44]. **E.** Clustering of genes showed consistent expression trend in both transcriptome and proteome data. Color as in Figure 3A.

Our proteomics (interactome, phosphoproteome, and ubiquitylome) analysis identified the significant enrichment of proteins related to mRNA splicing. Therefore, we performed the alternative splicing analyses of our transcriptome results. We observed 381 and 514 alternative splicing events in CoV2-NSP3- and CoV1-NSP3-expressing cells, respectively (Figure 7C). Among them, a large portion of altered splicing events was detected on the exon cassette and intron retention region, indicating that both NSP3 proteins alter host cell RNA processing.

Further, we performed FASP-based proteome analysis of NSP3-expressing cells in parallel. Differentially expressed proteins (DEPs) were calculated with thresholds of FDR < 0.05, and |L2FC| > 1 (Figure 7D). Among them, we found that a known target of CoV1-NSP3, TP53, was downregulated by both NSP3 proteins, and this downregulation was not CS-dependent [43]. Unfortunately, no significant GO terms or pathways were enriched for these DEPs. We compared our results with the proteome data after SARS-CoV-2 infection reported by Bojkova and colleagues [44] and highlighted overlapping proteins on the left of the heatmap (Figure 7D). A comparison of transcriptome and proteome data revealed that NF-*κ*B2 was upregulated at both levels, indicating a positive feedback regulation and highlighting the pivotal roles of NSP3 in NF-*κ*B signaling (Figure 7E).

### Multi-omics reveal cellular events regulated by NSP3 and the potential repurposing of drugs

We combined all of our significant candidates to systematic analysis CoV2-NSP3 regulated cellular events. We included known protein-protein interactions and regulation events to expand the connections of our candidates. Among the many cellular events we found, TP53 was one of the proteins that was extensively regulated by NSP3 (Figure 8A). We found interactors, activators, and inhibitors of TP53 in our multi-omics studies. In addition, these regulators were associated with other candidates, further expanding the diffusion network. Therefore, we concluded that NSP3 attacks p53 at multiple levels to circumvent the antiviral defense effects of p53 [43, 45]. Furthermore, the innate immune response, a surveillance system that SARS-CoV-2 needs to evade [46], was determined to be regulated by NSP3 expression (Figure S10). Besides regulating antiviral effects, NSP3 contributes to SARS-CoV-2-induced RNA alternative splicing [34]. We observed that proteins related to RNA splicing are regulated by NSP3 at post-translational levels (Figure 8B). For example, the phosphorylation of splicing factors and ubiquitination of heterogeneous nuclear ribonucleoproteins were changed in NSP3 expressing cells. Collectively, these data imply that NSP3 has diverse cellular effects in SARS-CoV-2 infection.

**Figure 8.**
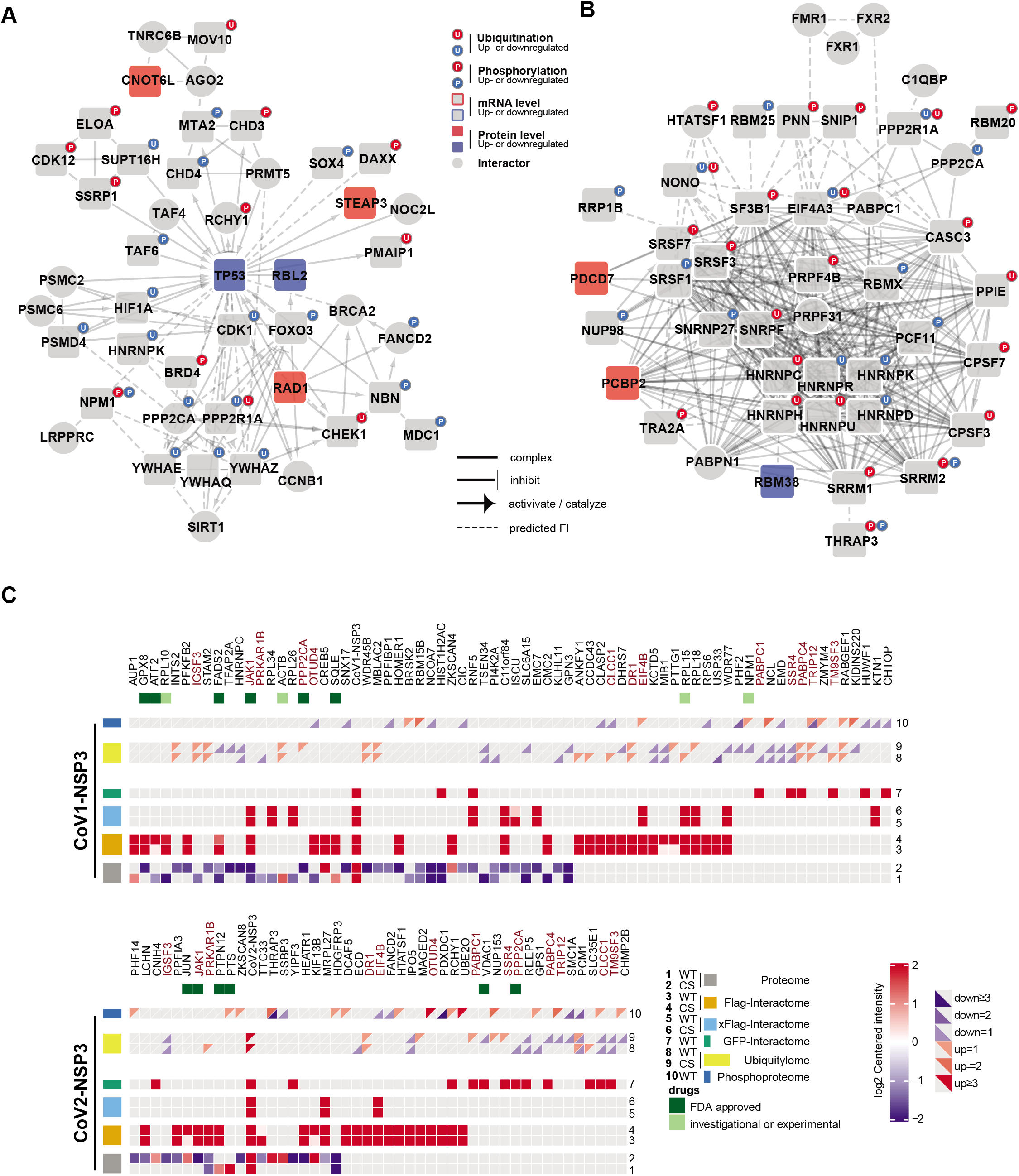
Regulation of cellular events by NSP3 and potential repurposing drugs for COVID-19. **A and B.** Regulation of p53 and alternative splicing by NSP3. Multi-omics data were combined to model the regulation of p53 (A) and alternative splicing (B) by NSP3. Network was created using Cytoscape with Reactome FI v7.2.3. FI represents functional interaction. **C.** Candidate proteins for drug repurposing. Candidate proteins that are regulated by NSP3 in at least two omics studies were shown. The rectangular is divided into two triangles to indicate upregulated or downregulated ubiquitination or phosphorylation events. Color as in Figure 3A. Protein with FDA-approved, or investigational drugs are labelled with green square.

Finally, we analyzed our data to identify proteins that were significantly enriched in at least two omics studies of NSP3 expressing cells (Figure 8C). In the meantime, we searched these candidate proteins against the drug database drugbank (https://go.drugbank.com/). In total, 13 proteins were regulated by two NSP3 proteins, and two (JAK1 and PPP2CA) out of them had nine FDA-approved drugs. Importantly, drugs that target JAK1 and PPP2CA are under investigation for COVID-19 [47–49]. Therefore, it would be interesting to investigate the antiviral effects of these candidate proteins with FDA-approved drugs.

## Discussion

As the largest polypeptide of SARS-CoV, NSP3 interacts with host cellular functions in a versatilely manner and plays a pivotal role in the viral life cycle. To meaningfully study the functions of NSP3, it needs to be expressed in its full-length form in mammalian cells. Recent reports demonstrated that PLpro alone is not sufficient to cleave the NSP1-NSP2 polypeptide, and the PLpro domain has a very similar interactome with NSP3 without the transmembrane region, suggesting that the localization and integrity of the NSP3 are important for its function and interaction with host cells [18, 21]. A previous study using deletions showed that CoV2-NSP3 has a general cytosolic localization, which is contradictory to our results [50]. We showed that CoV2-NSP3 has both ER and mitochondrial localizations, whereas CoV1-NSP3 localizes to the ER as reported [15]. Therefore, we believe that an interactome study based on NSP3 protein with proper localization is more reliable. By comparing of our results with a recently published interactome study using NSP3 deletions, we found that approximately 20% of interactors are shared between both studies. Importantly, we observed a strong association between NSP3 proteins and ribosomal protein in our study, which is lacking in the previous study.

Understanding the contributions of each SARS-CoV-2 protein to the functions of host cells is important to follow the pathogenesis of viral infection, and eventually develop antiviral compounds. Multi-omics methods, such as proteomics-based interaction and proteome studies have been proven to be useful tools for studying virus-host cell interactions [29, 44]. In this study, we combined the interactome, phosphoproteome, transcriptome, proteome and ubiquitylome to provide an integrative landscape of unique and shared functions of the NSP3 proteins. These parallel multi-omics datasets provide a rich resource for future mechanistic studies on how NSP3 regulates host cell functions in detail. Overall, we showed that both NSP3 proteins are connected to multiple fundamental cellular functions, including RNA metabolism, the immune response, cell cycle, translation, transcription, and others (Figure 9). These host cellular functions and signaling pathways are also largely affected by SARS-CoV infection [29, 44], indicating that NSP3 is one of the key polypeptides of SARS-CoVs in attacking host cell functions. Interestingly, multi-omics results showed that although both NSP3 proteins regulate similar cellular functions, they utilize different proteins to achieve these regulatory functions. For example, the CoV2-NSP3 prefers to associated with the mitochondrial translational machinery, whereas CoV1-NSP3 is linked to the cytosolic translational machinery. These results suggest a conserved cellular effect of SARS-CoVs NSP3 during the evolution.

**Figure 9.**
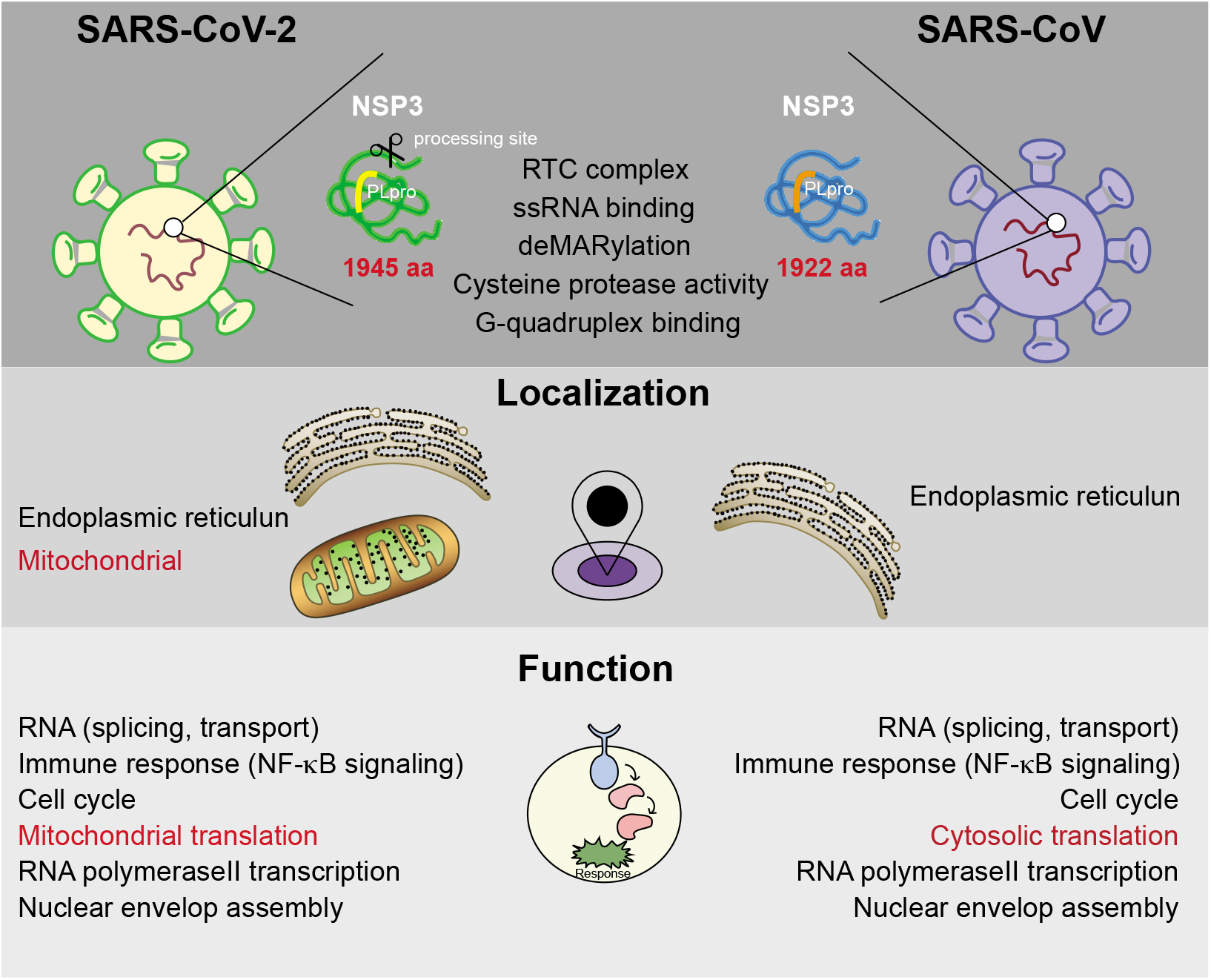
Multi-omics analysis of NSP3 functions. Schematic overview of NSP3 regulated cellular functions. Red indicates difference of two NSP3 proteins.

Previous studies on SARS-CoV have suggested that the combination of three transmembrane-localized NSPs (NSP3, NSP4, and NSP6) induces DMV formation, which together with other NSPs and host cell proteins, forms the RTC. Therefore, the membrane localization of NSP3 might be a key factor that the virus uses to select the organelles for RTC formation. In the present study, we found that unlike CoV1-NSP3, CoV2-NSP3 has two cellular localizations, N-terminal mitochondrial and C-terminal ER localizations. In addition, we showed that the N-terminal enrichment of CoV2-NSP3 extensively results in the immunoprecipitation of mitochondrial proteins. Furthermore, Wu and colleagues [51] showed that the SARS-CoV-2 RNA genome and sgRNAs are enriched in the host mitochondrial matrix using computational modeling. These results suggest that in SARS-CoV-2 infection, NSP3 might exploit both ER and mitochondria to initiate the RTC formation. In addition, we showed that proteins related to mRNA exporting and translation are enriched with both NSP3 proteins. In combination with the fact that the RTC is the location for RNA synthesis, we speculate that NSP3 couples RNA transport and translation together to rapidly synthesize viral proteins. However, we should note that the interaction between ribosomal proteins and NSP3 is not mediated by RNA, as we introduced universal nucleases to our cell lysis processes.

In summary, we expressed and studied the localization of NSP3 proteins. We identified a rich candidate resource for studying the functions of NSP3 proteins and provided a systematic analysis of how NSP3 contributes to the regulation of cellular function. Importantly, we showed that although both NSP3 proteins shared extensive similar functions, SARS-CoV-2 NSP3 is different from SARS-CoV NSP3 in its connection with mitochondrial proteins. Moreover, we provided a list of FDA-approved compounds that could serve as a starting point for drug repurposing with some already under investigation.

## Material and Methods

### Plasmids, Cell culture, Transfection, Immunoblotting and Immunoprecipitation

Full-length coding sequences of SARS-CoV-1 and SARS-CoV-2 NSP3 (FJ882960 and MN988669, without codon optimization) with earlier inserted stop codon were synthesized from IGEbio, China. PCR amplified products were subcloned into homemade lentivirus vector either with C terminal EGFP or N-terminal Flag and C-terminal EGFP tags. Mutagenesis PCR was performed to mutate the earlier stop codon to generate wild type NSP3 using Phanta Max Super-Fidelity DNA polymerase (P505-D2, Vazyme, China). The same method was used to create catalytic inactivated (CS mutated), glygly deficient (dGG) and domain exchanged NSP3 plasmids. UBE2O plasmid was described before [52] and TNIP cDNA was amplified from HeLa cDNA and cloned in to pCS2 vector with Flag tag. All batches of purified plasmids (Qiagen, 12165) were confirmed by DNA sequencing.

HEK293T cells from ATCC were cultured in Dulbecco’s Modified Eagle’s Medium (DMEM, Hyclone, SH30033.01) supplemented with 10% Fetal bovine serum (FBS, Hyclone, SV30160.03) and 1*×*Penicillin/Streptomycin (Hyclone, SV30010-10). HEK293T cells were tested mycoplasma free using MycoAlert Detection Kit (Lonza, LT07-318) before the whole experiments.

HEK293T cells were transfected using PEI (Polysciences, 24765-2) with a ratio of plasmids: PEI=1:3. Cells were harvest using whole cell lysate buffer (0.5% NP-40, 150 mM NaCl, 50 mM Tris pH 8.0, 10% Glycerol) with 1×cOmplete™ Protease Inhibitor Cocktail (11873580001, Sigma) 48 hr. post-transfection, following with a maximum speed centrifugation at 4°C. Supernatant were collected and protein concentration was measured using BCA assay kit (23225, Thermo). Equivalent amounts of proteins were boiled with 1*×*LDS loading buffer (NP0008, Thermo) for 5 min at 95°C and resolved by SDS-PAGE and immunoblotting. For immunoprecipitation, Flag-M2 agarose beads (A2220, Sigma) were used to enrich NSP3 and interactors. For phosphorylation validation, 1×PhosSTOP (4906837001, Sigma) was added to lysis buffer, and immunoprecipitation was performed either with Flag agarose beads for 2 hr. or with 1 μg UBE2O antibody (A301-873A, Bethyl) overnight following with 15 μL prewashed protein A/G beads (1:1 mixture, 1614013 and 1614023, Biorad) incubation for another 1.5 hr. Bound proteins were boiled with 1*×*LDS loading buffer for 5 min at 95°C after 3 washes with lysis buffer. Antibodies used in these experiments included Flag (F1804, Sigma), EGFP (50430-2-AP, Proteintech), NSP3 (ab181602, Abcam), UBE2O (A301-873A, Bethyl), pSerine (gtx26639, GeneTex), pThreonine (9386, Cell Signaling), TAB2 (A302-759A, Bethyl), TAB3 (A302-208A, Bethyl), RPL5 (A303-933A, Bethyl), RPS6 (A300-556A, Bethyl), MRPL13 (16241-1-AP, Proteintech), MRPL23 (11706-1-AP, Proteintech), MRPL44 (16394-1-AP, Proteintech), USP34 (A300-824A, Bethyl), DHX38 (A300-858A-T, Bethyl), IPO4 (11679-1-AP, Proteintech), IPO11(14403-1-AP, Proteintech), NUP85 (19370-1-AP, Proteintech) and GAPDH-HRP (KC-5G4, KANGCHENG, China).

### Immunofluorescence

Cells grown on 0.1% gelatin coated coverslips were transfected with indicated plasmid for 36 hr. 4% paraformaldehyde was used to fix the cells at room temperature for 30 min followed with quenching with 2 mg/mL glycine. Cells were then permeabilized with 0.2% Triton X-100 and blocked with 10% FBS. Indicated first antibody and second antibody were added for overnight at 4℃ and 1 hr. at room temperature, respectively. Cells were mounted with ProLong Gold Antifade Reagent with DAPI (Cell Signaling, 8961S). Images were taken and processed using LSM800 and LSM900 from Zeiss. Antibodies used for the experiment are: CANX (2679, Cell Signaling), TOMM20 (ab56783, Abcam), Flag (F1804 and F7425, Sigma), NSP3 (ab181602, Abcam).

### Fluorescence-activated Cell Sorting (FACS)

Cell cycle and mitochondrial potential experiments were performed using FACS. For cell cycle analysis, approximately 1mL of 1*×*10^6^/mL cells transfected with indicated plasmids were harvest and fixed with 70% ethanol overnight. 0.1% Triton-X100 was used to permeabilize cells before the staining with propidium iodide for 15 min at room temperature. Cells were analyzed with PE channel using Beckman CytoFLEX.

### Omics sample preparation

All omics experiments were performed in biological replicates.

### Interactome sample preparation

HEK293T cell transfected with indicated plasmids were lysed using whole cell lysis buffer (50 mM Tris pH 8.0, 150 mM NaCl, 10% Glycerol, 0.5% NP40) with fresh added 1*×*Complete Protease inhibitors and Universal Nuclease (1:5000, 88702, Thermo). For cross-linking IP-MS with RIME method, transfected cells were incubated with 2 mM discuccinimidyl glutarate (sc-285455A, SantaCruz) for 20 min at RT first, then replaced with 1% formaldehyde (F8775, Sigma) for 10 min at RT, followed with quenching with 0.1M glycine for another 10 min at RT. Cells were incubated for 2 hr. at 4°C on a rotation wheel. Soluble cell lysates were collected after maximum speed centrifugation at 4°C for 15 min. 1 mg of cell lysates were incubated with Flag-M2 (M8823, Sigma) or GFP (gtma, Chromotek) magnetic agarose for 1.5 hr. On-bead digestion was performed to digest immunoprecipitated protein to peptides for MS measurements [53]. In short, beads after incubation were washed 3 times with cell lysis buffer and 1 time with PBS. After completely removal of PBS, beads were incubated with 100 μL of elution buffer (2 M urea, 10 mM DTT and 50 mM Tris pH 8.5) for 20 min. Afterwards, IAA was added to a final concentration of 50 mM for 10 min in the dark, following with 250 ng of trypsin (V5280, Promega) partially digestion for 2 hr. After incubation, the supernatant was collected into a separate tube. The beads were then incubated with 100 μL of elution buffer for another 5 min, and the supernatant was collected in the same tube. All these steps were performed at RT in a thermoshaker C at 1500 rpm. Combined elutes were digested with additional 200 ng of trypsin overnight at room temperature. Finally, tryptic peptides were acidified to pH < 2 by adding 10 μL of 10% TFA (Sigma, 1002641000).

### Phosphoproteome sample preparation

HEK293T cells transfected with indicated plasmids were lysed using SDC (D6750, Sigma) buffer (4% SDS, 100 mM Tris pH 8.5) and phosphoproteome sample preparation was performed according to previous report [33]. In brief, 200 μg of cell lysates were reduced and alkylated with Tris (2-carboxyethyl) phosphine hydrochloride (TCEP, 75268, Sigma) and 2-chloroacetamide (CAA, C0267, Sigma) at 45°C for 5 min, following with overnight digestion with Lys-C (129-02541, Wako Chemicals) and trypsin (T6567, Sigma). TiO_2_ beads (5010-21315, GL Sciences) were used to enrich phosphorylated peptide and homemade C8 (66882-U, Sigma) stageTip was used to trap the TiO_2_ beads for washes. Eluted phosphorylated peptides were loaded into homemade SDB-RPS (66886-U, Sigma) stageTip for desalting.

### Ubiquitylome sample preparation

HEK293T cell transfected with indicated plasmids were lysed using buffer containing 8M urea, 50 mM Tris-HCl, pH 8, 150 mM NaCl, 1 mM EDTA, 1 mM chloroacetamide and 1*×*Complete Protease inhibitors. K-*ε*-GG ubiquitin Remnant Motif enrichment kit from Cell signaling (5562, Cell Signaling) was used to enrich K-*ε*-GG peptides. The whole process was performed according to the manufacturer’s instruction except in the steps of digestion and peptide desalting. We used Lys-C to digest 3 mg protein in 8M urea buffer for 3 hr. at 37°C first, and then diluted to 2M urea and trypsinization overnight at 37°C. For peptide desalting, we used the Blond Elute LRC-C18 200 mg column from Agilent (12113024), followed with lyophilization for 2 days.

### Proteome sample preparation

HEK293T cell transfected with indicated plasmids were lysed with 8 M urea with 10 mM DTT, 0.1 M pH 8.5 Tris buffer. Soluble cell lysates were collected after 10 cycles of sonication for 30s followed by 30s at 4°C using a Diagenode Biorupter Pico Sonicator, followed with maximum speed centrifugation at room temperature for 15 min. Whole proteome peptides were prepared using the FASP protocol as described before [53]. In brief, 50 μg of soluble lysates was added onto 30 kDa cut-off filter (MRCF0R030, Millipore) and centrifuged at 11000 rpm at 20°C for 15 min. 50 mM iodoacetamide (IAA, Sigma, I1149) in urea buffer was used to alkylate proteins at 20°C for 15 min. After 3 washes with urea lysis buffer and 3 washes with 50 mM ammonium biocarbonate (ABC) buffer, 500 ng of trypsin in 50 μL 50 mM ABC buffer was used to digest proteins in a wet chamber overnight at 37°C. Peptides was extracted by 50 mM ABC buffer and acidified to pH < 2 by adding 10 μL of 10% TFA.

### RNA sequencing

HEK293T cell transfected with indicated plasmids were harvested and total RNA was isolated using ipureTRizol kit according to manufacturer’s instruction (K417, IGEbio). Sequencing libraries were generated using Next UltraTM RNA Library Prep Kit for Illumina (E7760, NEB) following manufacturer’s recommendations. In brief, mRNA was captured using mRNA capture beads, followed with fragmentation and cDNA synthesize using random hexamers. DNA clean beads was used to purified dsDNA after the second strand synthesis, followed with end repairing, A tailing, adapter ligation, PCR amplification and library purification. Sequencing was done by Illumina NovaSeq platform (IGEbio, China).

### Mass spectrometry measurements

Desalted peptides were separated and analyzed with an Easy-nLC 1200 connected online to Fusion Lumos or Fusion Eclipse (cross-linking experiment) mass spectrometer equipped with FAIMS pro using different gradient of buffer B (80% acetonitrile and 0.1% Formic acid). For interactome study, a gradient of total 140 min of 100 min 2% to 22%, 20 min 22% to 28%, 12 min 28% to 36% and 8 min 100% of buffer B was used; for proteome and ubiquitylome analysis, a gradient of total 240 min of 1 min of 2%, 10 min 2% to 7%, 200 min 7% to 28%, 15 min 28% to 36%, 5 min 36% to 60% and 7 min 95% of buffer B was used; for phosphoproteome analysis, a gradient of total 120 min of 80 min 2% to 22%, 20 min 22% to 28%, 12 min 28% to 40% and 8 min 95% of buffer B was used. Data dependent analysis was used as data acquisition mode. Detailed information about the gradient and the setting of the MS can be found in the raw files. Raw data were first transformed using FAIMS MzXML Generator and then analyzed using MaxQuant version 1.6.17.0 [54] search against Human Fasta database with CoV1-NSP3 and CoV2-NSP3 protein sequences. Label free quantification and match between runs functions were enabled for all analyses. For phosphoproteome and ubiquitylome analyses, pSTY and Gly-Gly (K, not C-term) in variable modifications were added.

### Data Analysis

PCA for all omics studies were performed using top 1000 variable events with maximum standard deviation of NSP3-expressing cells. PC1-PC2 scatterplot exhibits the magnitude of the difference between samples.

### Proteome and Interactome data analysis

The MaxQuant output files “proteinGroup” was used for the subsequent analysis of proteome and interactome. We filtered out the protein that labelled as reverse or potential contaminant, and analyzed the non-crosslinking interactome and proteome data using DEP [55]. The significant thresholds were L2FC > 2, FDR < 0.001 for interactome, and |L2FC| > 1, FDR < 0.05 for proteome analyses. For crosslinking interactome analysis, we used an alternative method described before, which used a variable filter combining log2 fold-change (enrichment) and adjusted *P* value [56]. Briefly, to determine enriched interactors, we used a cutoff lines with the function y > *c*/(x - x0) on scatters where x is the log2 fold-change and y is adjusted P value (*c* = curvature, x_0_ = minimum fold change). The distribution of log2 fold-changes between NSP3s overexpression and EGFP control were fitted to a Gaussian curve using least squares fitting (excluding outliers) to determine the standard deviation σ. We set a stringent fold-change cutoff x_0_ to 2 σ while selected a relatively loose *c* to get a higher positive coverage rate. Then we verify the overlap between crosslinking IP-MS and non-crosslinking IP-MS under selected curvature *c*.

Protein-Protein Interaction (PPI) network and pathway annotation of interactors was constructed by Cytoscape plugin Reactome FI v7.2.3 [57]. Interactors of NSP3 were divided into 3 groups (CoV1, CoV2 and both) and enriched GO terms were shown using clusterProfiler package [58]. Circos plots was performed using circlize package [59].

### Phosphoproteome data analysis

The MaxQuant output files designated “Phospho(STY)sites” was used for testing intensity differences of phosphosites between control and NSP3 expressing cells. We performed Student’s t test as two-sample-test and one-way analysis of variance (ANOVA) test following Tukey’s significant honest difference post-hoc test (THSD) as multiple-sample-test. *P* values were adjusted by Benjamini & Hochberg (BH) method. We defined peptide with p-adjusted < 0.01, |L2FC| > 2 and localization possibility > 0.7 as significant enriched peptide.

To study change of kinase activities, we ranked phosphosites by L2FC and performed Kinase-Substrate Enrichment Analysis (KSEA) using KSEAapp package [60] in combination with a reported kinase-substrate relationship [61]. We filtered out the kinases of which matching sites was less than 3 or Z-score is lower than 1 and created Human Kinome Tree illustration using KinMap [36]. The targeting drugs that regulate activity changed kinases were extracted from drugbank (https://go.drugbank.com/).

### Ubiquitylome data analysis

The ubiquitylome data was analyzed as described for phosphoproteome. Here, we defined peptide with adjusted p < 0.001, |L2FC| > 2 and localization possibility > 0.7 as significant enriched peptide.

### Transcriptome data analysis

Raw reads were aligned to the human genome (hg38) using HISAT2 [62]. Next, we assembled reads to transcripts and quantified the read counts of each genes utilizing StringTie [63]. Lowly expressed genes were filtered out using a CPM (counts per million) threshold. We kept genes that CPM is greater than 1 in at least 2 samples and analyzed the differential expression gene using edgeR [64]. Top 1000 upregulated and 1000 downregulated genes were ordered by L2FC as the gene list input of GSEA analysis. Differentially expressed genes (DEGs) between control and NSP3 expressing cells were characterized for each sample with thresholds of |L2FC| > 1, FDR < 0.05.

### Motif Enrichment

The DEGs were divided into 2 gene sets according to their expression change status (cluster 1&2 for upregulated genes and cluster 3&4 for downregulated genes) as represented in Figure 7A. Then, the putative TF-targeting regulons of these 2 gene sets were identified by employing RcisTarget [65]. We choose human motif collection v9 as TF annotation and the motif-rankings in 10 kbp around transcription start sites (TSS) of hg38 genome as region databases. The motifs with most target genes as well as a normalized enrichment score (NES) lager than 3 were determined as significant enriched regulons.

### Alternative Splicing Analysis

We performed alternative splicing (AS) analysis based on transcription raw reads with vast-tools V2.5.1 [66] in combination with vastdb hs2.23.06.20. To determine significant AS events, we kept the AS event which have more than 10 reads in at less two samples and applied a further filter by setting a noB3 parameter to AltEx and a 0.05 binomial-test p_IR parameter to IR in vast-tools’ diff module. Significant AS events were defined according to the following requirements: change was greater than 10 dPSI/dPIR, minimal difference value where *P* > 0.95 was greater than 0.

### Gene Ontology and Gene Set Enrichment Analysis

GO (Gene ontology) and Gene Set Enrichment Analysis (GSEA) were implemented based on clusterProfiler package. C2 (curated gene sets), C5 (GO gene sets) from MSigDbv7.2 [41] and the pathway gene sets from Reactome Pathway Database [30] were chosen for GSEA analysis. Significant terms were chosen by the threshold that *P* value < 0.05, FDR (false-discovery rate) < 0.25 and |NES| > 1. Most significant terms were selected to plot heatmap.

The Over Representation Analysis (ORA) was based on the hypergeometric distribution and *P* value was adjusted by BH method. Gene ontology analysis of Biological Processes (BP), Cellular Component (CC), Molecular Function (MF) were performed using enrichR in clusterProfiler package. Reactome pathway analysis was performed using enrichPathway function in ReactomePA packages [8]. Gene sets large than 500 or less than 15 genes were excluded from ORA. Significant GO terms were enriched by a threshold of adjusted *P* value < 0.05, q-value < 0.25 and |NES| > 1.

## Ethical statement

Not applicable

## Code availability

Not applicable

## CRediT author statement

**Shi Ruona:** Conceptualization, Investigation, Validation, Visualization **Feng Zhenhuan:** Conceptualization, Formal analysis, Data Curation, Visualization **Zhang Xiaofei:** Conceptualization, Supervision, Writing-Original Draft, Writing-Review & Editing, Funding acquisition.

## Acknowledgements

We like to thank Bioland Laboratory, Guangzhou, China for the generous startup package. Work in Zhang lab is supported by the Guangdong Science and Technology Project (2018B030306047, 2020B1212060052), National Natural Science Foundation (31770889, 31801180), Guangzhou Science and Technology Project (201904010469) and Guangzhou Regenerative Medicine and Health Guangdong Laboratory project (2018GZR110104003).

We apologize to all colleagues whose work were unable to be cited and discussed due to space constraints. We thank all colleagues whose excellent work helped us to understand and tackle the COVID-19. We like to thank Prof Duanqing Pei from Westlake university, China, for valuable suggestions. We like to thank all member in Zhang lab for fruitful discussions, Liman Guo from Proteomics & Metabolomics Centre of Bioland Laboratory for excellent mass-spectrometry measurements assistance, Yu Fu from Microscope & Imaging Centre of Bioland Laboratory for confocal imaging and Chan Rong from CCLA of Bioland Laboratory for FACS assistance.

## Supplementary material

### Supplementary Figure legends

**Figure S1.**
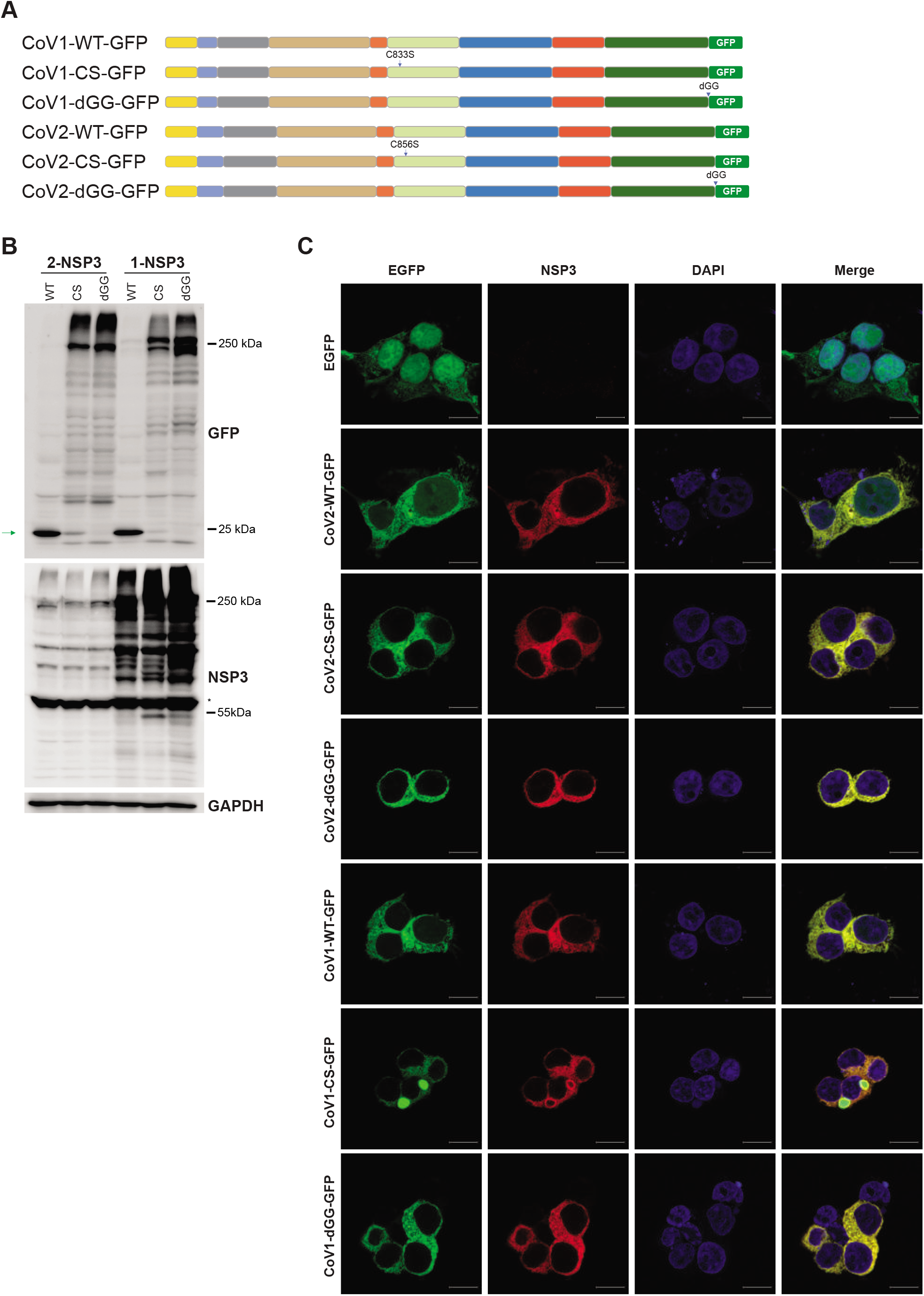
Expression and subcellular localization of NSP3 proteins with C-terminal tag. **A.** Schematic illustration of plasmids used in Figure 1B, Figure S1B and S1C. **B.** NSP3 cleaves itself at the C-terminus. HEK293T cells transfected with CoVs-NSP3-EGFP plasmids were lysed and EGFP and NSP3 antibodies were used after immunoblotting. EGFP was released in WT, but not in CS- or dGG-mutated NSP3 samples. Green arrow indicates released EGFP, whereas asterisk indicates non-specific signal. **C.** Validation of the specificity of CoV-1 NSP3 antibody. HEK293T cells transfected with indicated plasmids were fixed 48 hr. post-transfection. EGFP and NSP3 antibodies were used to detect the localization of NSP3 proteins.

**Figure S2.**
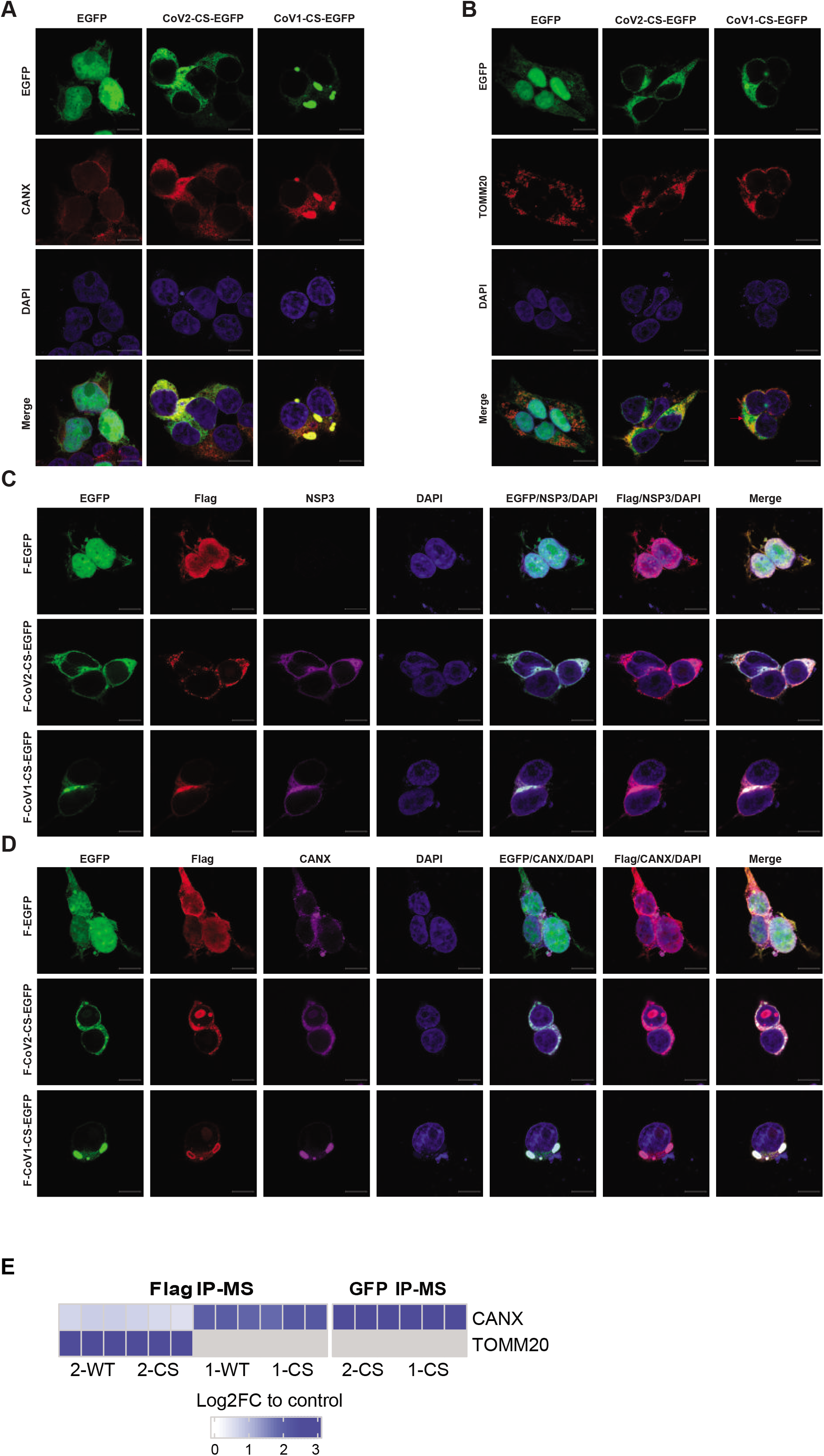
NSP3 proteins have preferable cellular localization. **A** and **B.** Cellular localization of NSP3 protein detected by EGFP antibody. HEK293T cells transfected with indicated plasmids were fixed 48 hr. post-transfection. EGFP fluorescence was used to detect the localization of CS-NSP3-EGFP protein. CANX and TOMM20 antibodies (red) were used to serve as markers for the ER (**A**) and mitochondria (**B**). Red arrow indicates that the large vesicle-like structure of CoV2-NSP3 was not co-localized with TOMM20. **C.** CoV2-NSP3 has two cellular localizations. Immunostaining was performed as in Figure S2A. Merged signals show that GFP and NSP3, but not NSP3 and Flag, are co-localized well. **D.** Flag-CoV1-NSP3 prefers to co-localize with CANX. Immunostaining was performed as in Figure S2A. Flag and CANX antibodies were used to visualize the co-localization between CoV1-NSP3 and the ER marker CANX. **E.** Interaction preference of NSP3 protein and CANX/TOMM20. Interactome studies of NSP3 based on Flag and EGFP IP-MS were analyzed to show the binding preference between NSP3 proteins and CANX/TOMM20.

**Figure S3.**
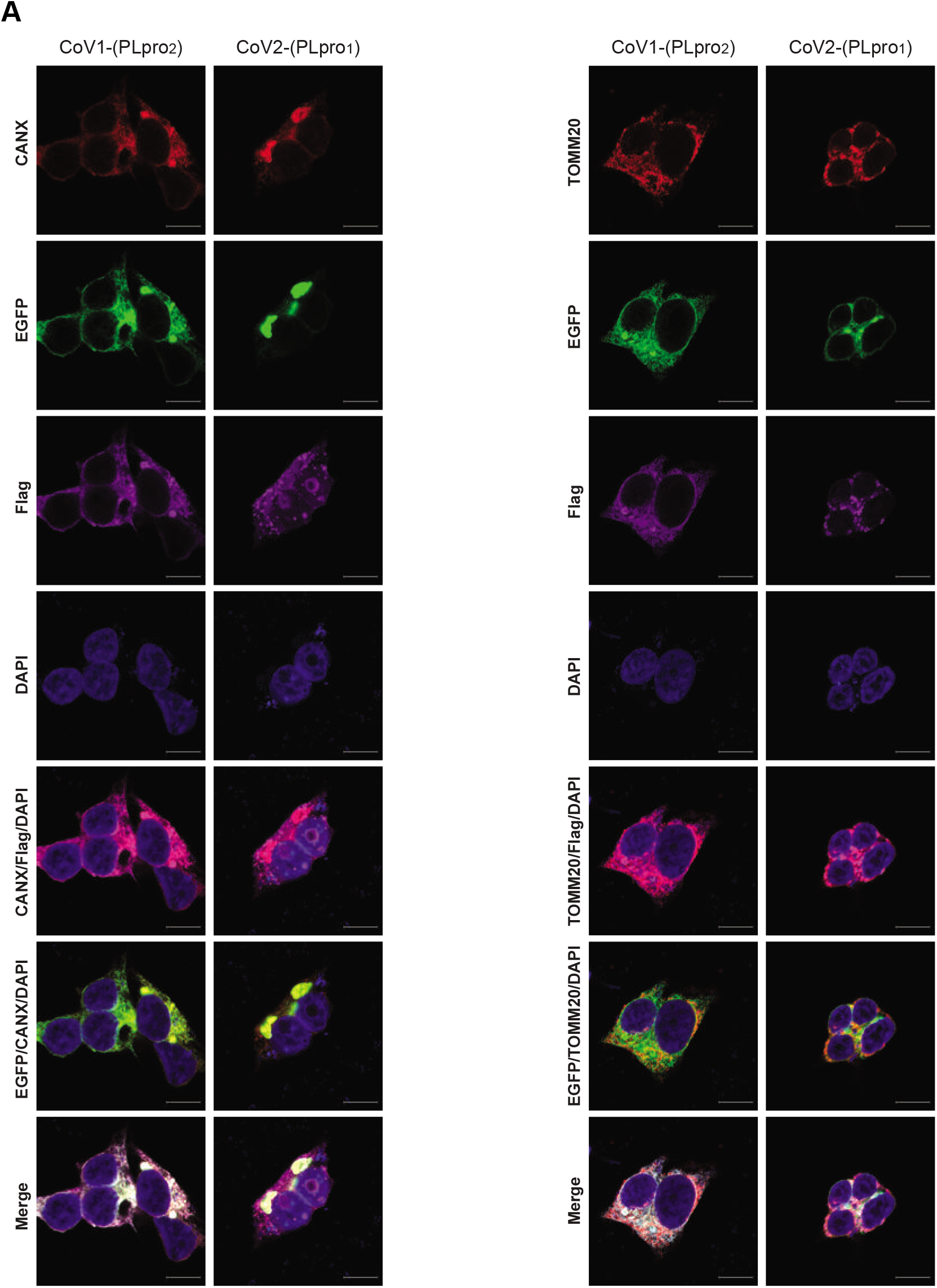
PLpro domain of Cov1-NSP3 is responsible for formation of the large vesicle-like structures. **A.** CoV2-NSP3 fused with PLpro domain of CoV1-NSP3 induces large-vesicle structures. Immunostaining was performed as in Figure S2A.

**Figure S4.**
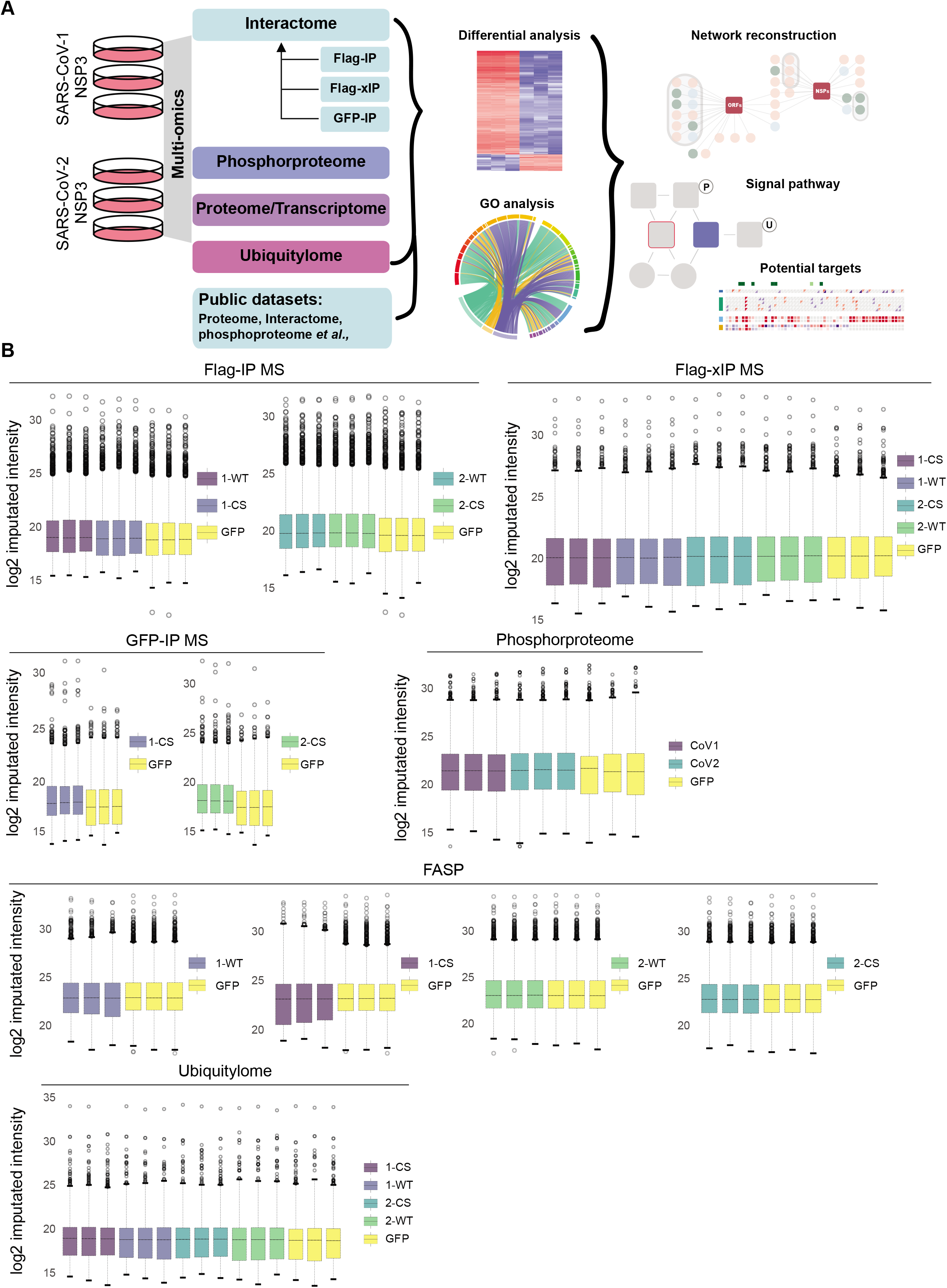
Workflow and quality control of multi-omics study. **A.** Schematic overview of the workflow to study the function of NSP3. **B.** Boxplots of log2 imputed intensity for all proteomics studies.

**Figure S5.**
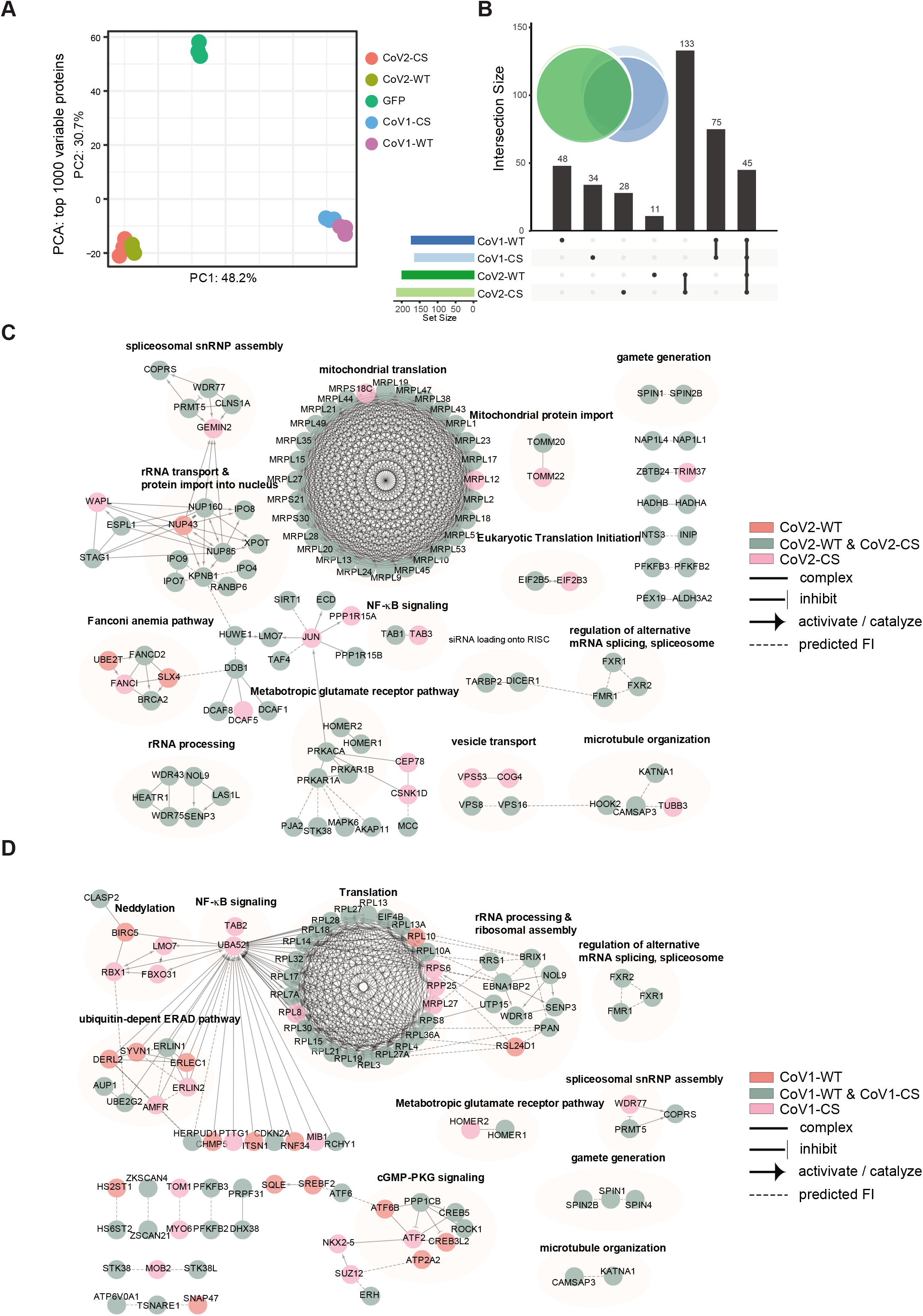
Protein-protein interaction network of NSP3 interactomes. **A.** PCA analysis of top 1000 variable proteins of NSP3 interactomes. Top 1000 variable proteins with highest standard deviation were used as input for PCA analysis. **B.** Upset and venn diagrams showing the overlap of NSP3 interactors. Significant interactors for WT and CS of NSP3 proteins were used as input. **C and D.** Protein-protein interaction network of CoV2-NSP3 (**C**) and CoV1-NSP3 (**D**) interactomes. Significant interactors of NSP3 were clustered by gene annotation and protein-protein interaction network was constructed using Cytoscape with Reactome FI v7.2.3. FI represents functional interaction.

**Figure S6.**
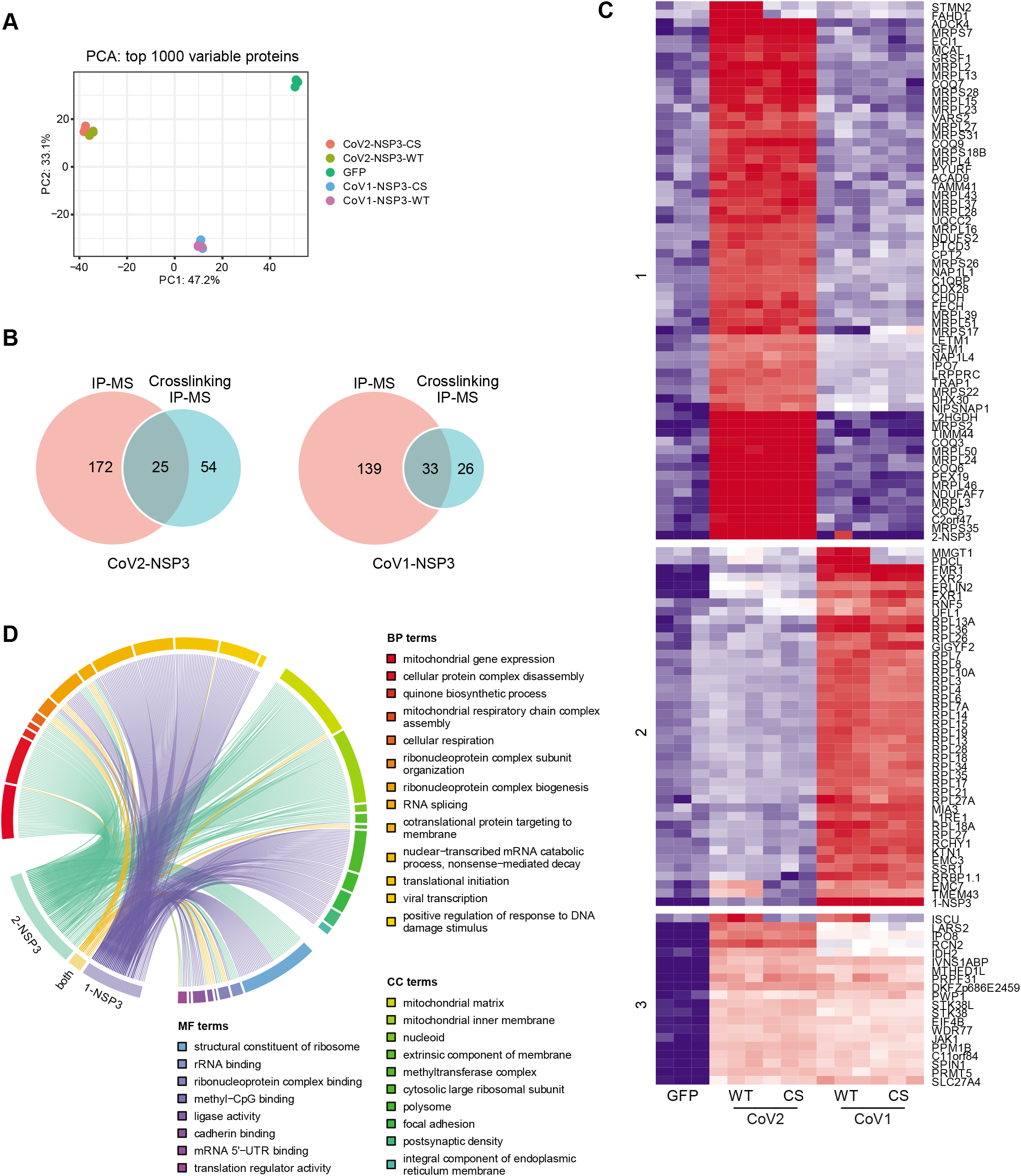
Crosslinking IP-MS to identify NSP3 interactome. **A.** PCA analysis of top 1000 variable proteins of NSP3 interactomes. Top 1000 variable proteins of each condition were used as input for PCA analysis. **B.** Venn diagram showing the overlap of NSP3 interactors identified in crosslinking and non-crosslinking experiments. **C.** Hierarchical clustering of significant interactors of NSP3 proteins using cross-linking IP-MS. Analysis and color as Figure 3A. **D.** Representative enriched GO of identified interactors. Analysis as Figure 3B.

**Figure S7.**
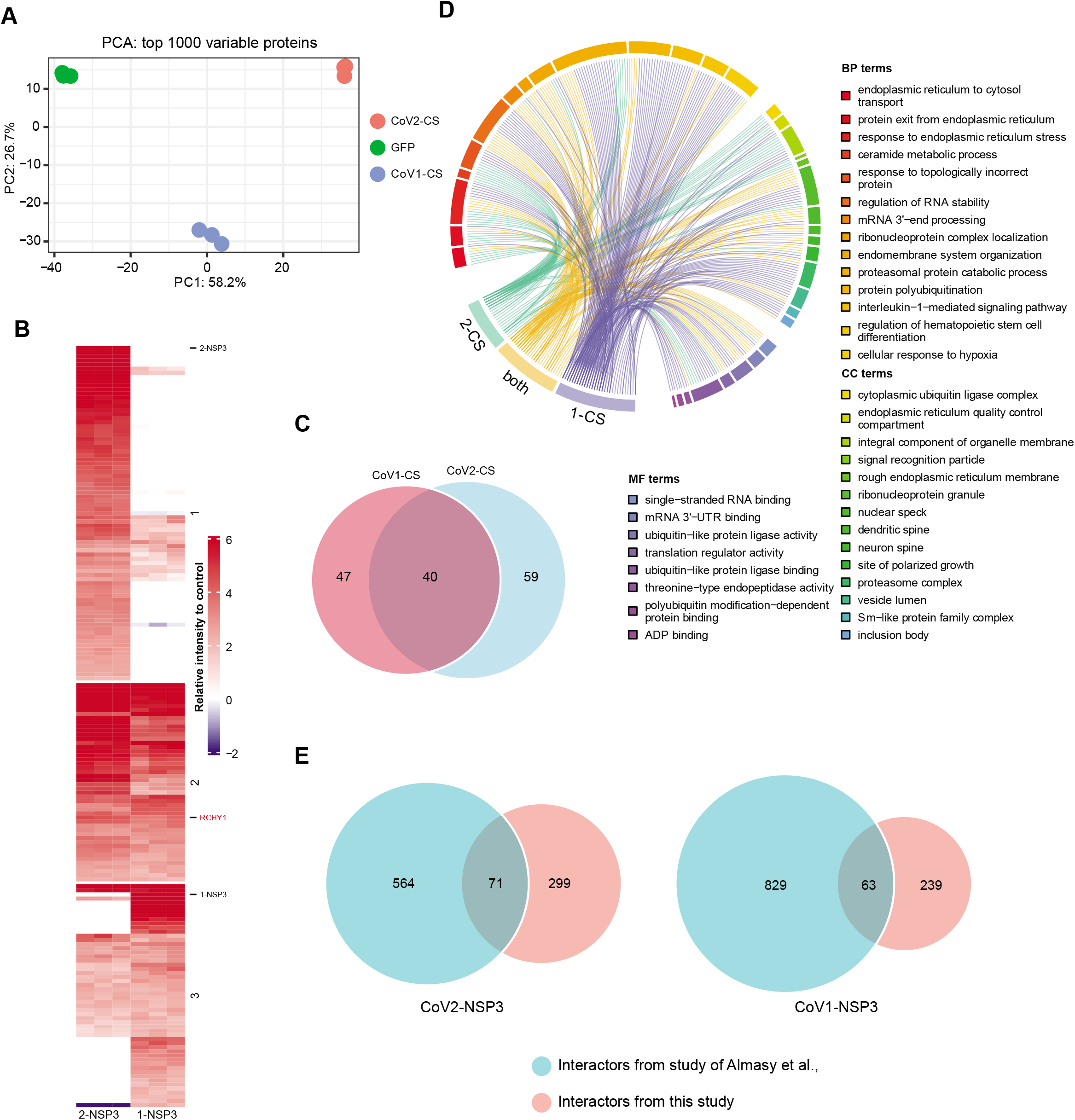
NSP3 interactome identified by C-terminal EGFP-IP. **A.** PCA analysis of top 1000 variable proteins of NSP3-EGFP interactomes. Top 1000 variable proteins with highest standard deviation were used as input for PCA analysis. **B.** Hierarchical clustering of significant interactors of NSP3 proteins. The relative intensity to control samples of significant interactors was compared between CoV1-NSP3 and CoV2-NSP3. **C.** Venn diagram showing the overlap of CoV1-NSP3 and CoV2-NSP3 interactomes. **D.** Representative enriched GO of identified interactors. Analysis as Figure 3B. **E.** Venn diagram showing the overlap of NSP3 interactors identified in present and previous studies.

**Figure S8.**
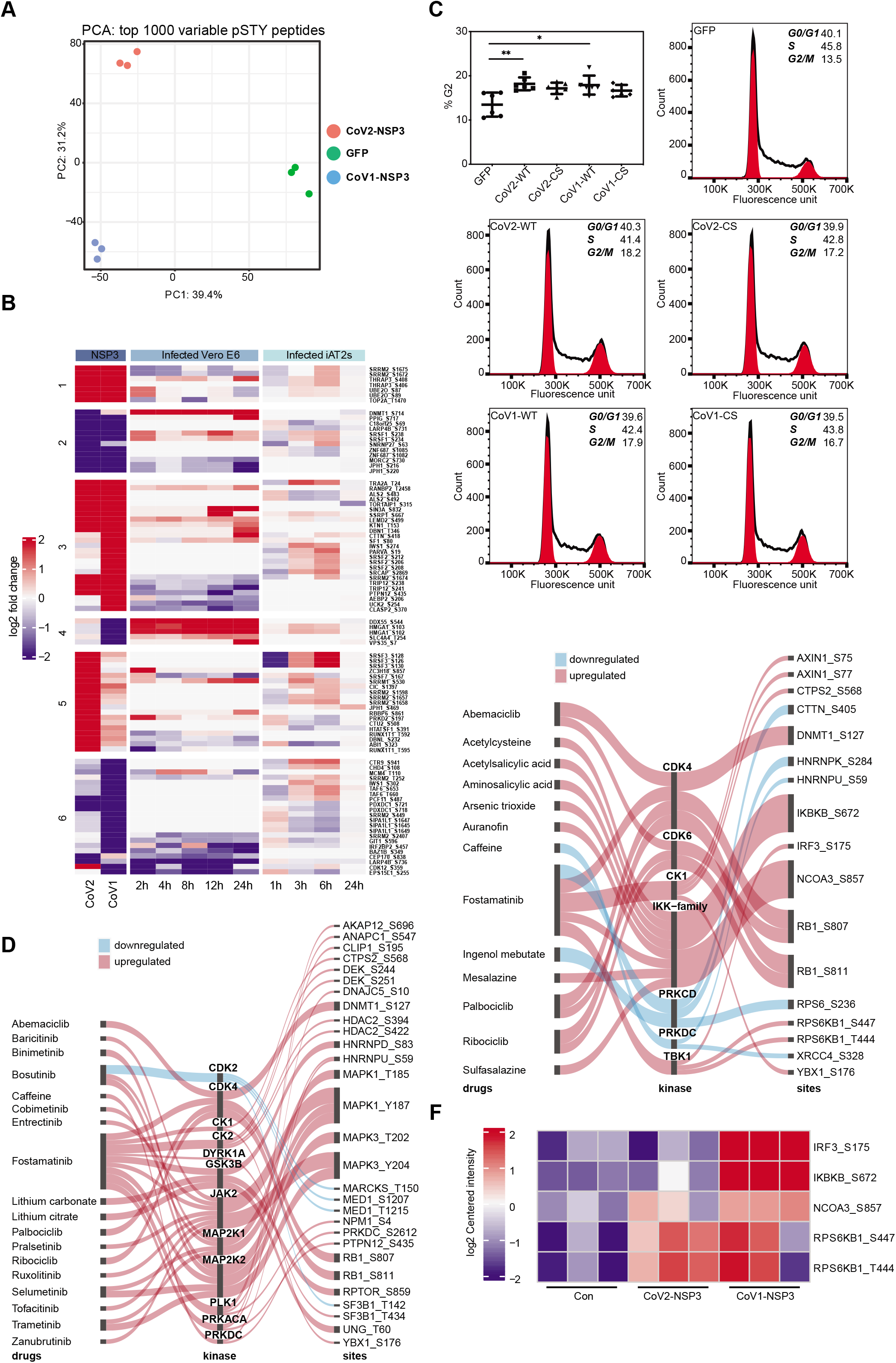
Phosphoproteome related analyses. **A.** PCA analysis of top 1000 variable phosphorylated peptides of NSP3 expressing cells. Top 1000 variable phosphorylated peptides with standard deviation were used as input for PCA analysis. **B.** Hierarchical clustering of DEPPs which were also found in SARS-CoV-2 infection studies. Two reported phosphoproteome data were used as inputs to identify consistent phosphorylated peptides. Analysis and color as Figure 3A. **C.** Cell cycle analysis of NSP3 expressing cells. Cells transfected with indicated plasmids were harvest for FACS analysis using propidium iodide. Significant differences compared to control cells were calculated using student t-tests. The graph showed mean ± SD, n=3, * indicates p < 0.05, ** indicates p < 0.01. One representative experiment was shown. **D and E.** Kinases inhibitors mapped to kinases whose activity mapped to DEPPs in CoV2-NSP3 **(D)** and CoV1-NSP3 **(E)** expressing cells. Red indicates upregulated activity, whereas blue indicates downregulated activity. **F.** Hierarchical clustering of DEPPs that regulated by IKK-family and TBK1. Analysis and color as Figure 3A.

**Figure S9.**
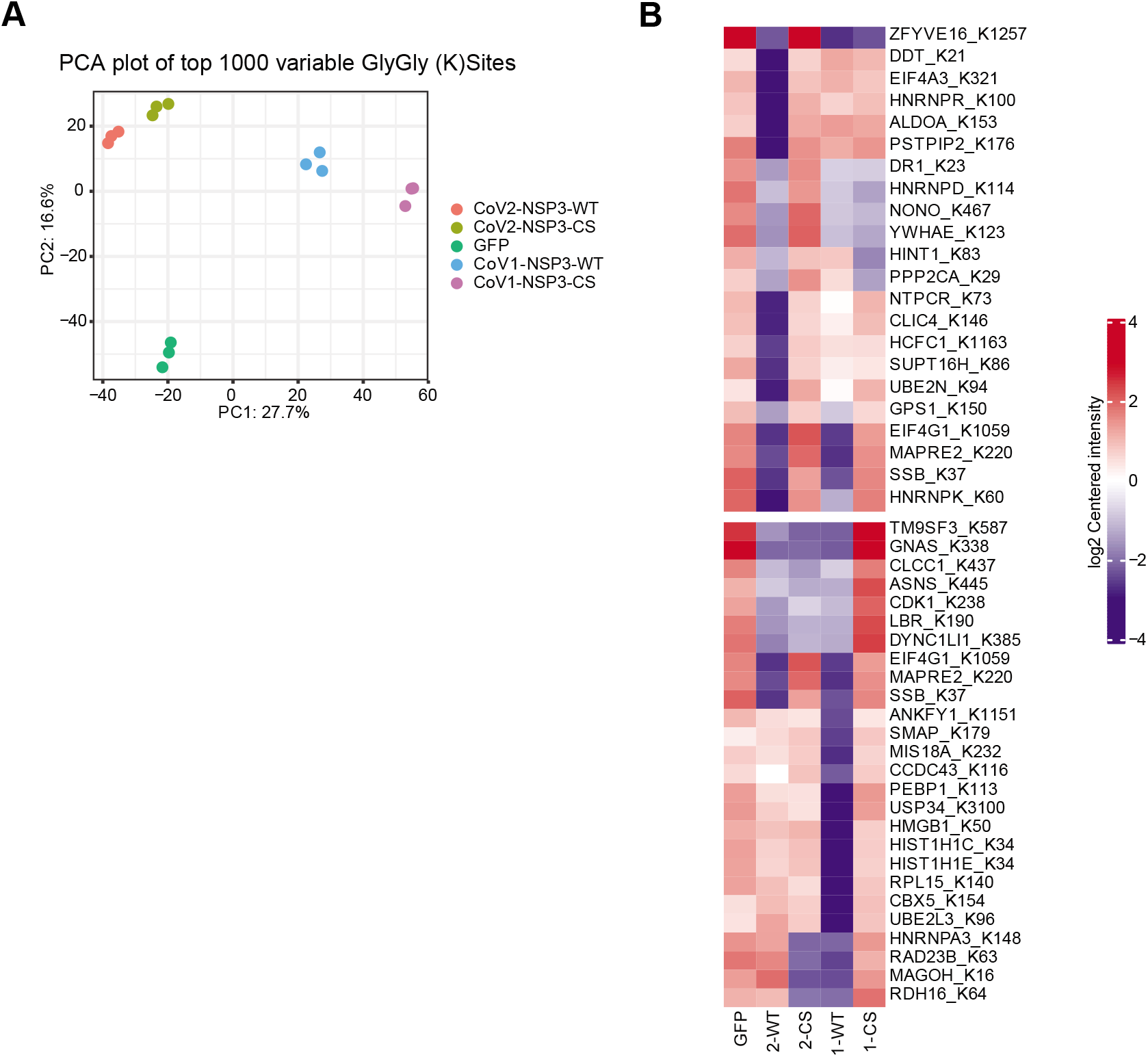
Ubiquitylome related analyses. **A.** PCA analysis of top 1000 variable ubiquitinated peptides of NSP3 expressing cells. Top 1000 variable ubiquitinated peptides with highest standard deviation were used as input for PCA analysis. **B.** Deubiquitinase activity dependent ubiquitylome of NSP3 protein. Analysis and color as in Figure 3A.

**Figure S10.**
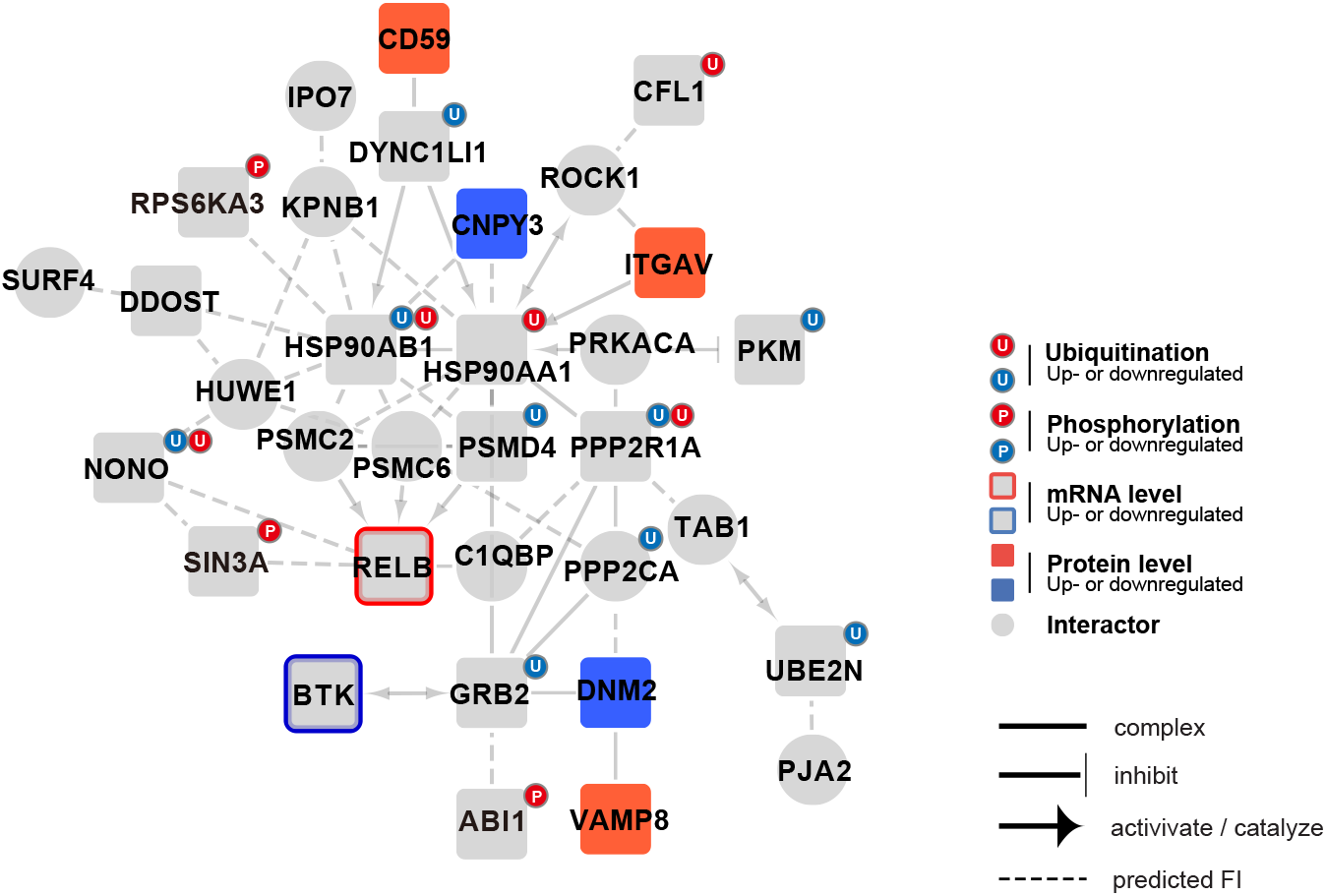
Regulation of innate response by NSP3. Analysis as Figure 8A.

